# Large-scale genomic analysis reveals the distribution and diversity of Type VI Secretion Systems in *Escherichia coli*

**DOI:** 10.1101/2025.01.23.634537

**Authors:** Kristina Nesporova, Sander K. Govers

**Affiliations:** Department of Biology, KU Leuven, 3001 Leuven, Belgium

## Abstract

The type VI secretion system (T6SS) is a versatile nanomachine that injects effectors into target cells, playing a role in bacterial competition and virulence. While widespread in Gram-negative bacteria, T6SS prevalence varies across species and strains, and its distribution in *Escherichia coli* remains underexplored despite *E. coli* being an important intestinal (IPEC) and extraintestinal (ExPEC) pathogen. Here, we examined the prevalence and distribution of T6SS subclasses (T6SS^i^) across 131610 *E. coli* genomes, which we annotated for clinical relevance. T6SS genes were identified while focusing on the three subclasses of T6SS^i^ gene clusters known to be present in *E. coli:* T6SS^i1^, T6SS^i2^, and T6SS^i4b^. Across phylogenetic groups, T6SS^i1^ was broadly present, while T6SS^i2^ showed associations with B1, B2, and G, and T6SS^i4b^ was rare. T6SS^i1^ was primarily associated with IPEC, and T6SS^i2^ with ExPEC. Even clearer patterns emerged at the sequence type (ST) level or when strains were further categorized into pathovars. For example, both the most dominant ExPEC and IPEC ST (ST131 and ST11, respectively) displayed niche-specific T6SS trends, with non-complete T6SSs being more associated with humans. In a final step, we evaluated the co-occurrence of T6SSs with other virulence-associated genes (VAGs) and multidrug resistance (MDR). This analysis confirmed the association of T6SS^i1^ and T6SS^i2^ with IPEC- and ExPEC-associated VAGs, respectively, and revealed a negative correlation between complete T6SS^i^ subclasses and MDR. Together, our work provides a comprehensive overview of the diversity of T6SSs across *E. coli*, shedding more light on their potential contribution to pathogenicity in this species.

**Importance:** Our study represents a large-scale analysis of T6SSs across one of the most comprehensive collections of *E. coli* genomes to date. In doing so, we updated several misconceptions on T6SSs distribution and other genomic properties of *E. coli* strains, which originated from smaller scale studies and were subsequently extrapolated in the literature. This includes the prevalence and distribution of T6SS^i^ subclasses across phylogenetic groups (e.g., T6SS ^i2^ is not prevalent in phylogroup D), the association of specific virulence factors with IPEC and/or ExPEC (e.g., hemolysin A is more often associated with IPEC and not a typical ExPEC characteristic), and characteristics of pathogenic STs (e.g., ST131 displays distinct genomic properties based on its environmental niche). As such, this study not only advances our understanding of T6SS in *E. coli*, but also serves as a valuable resource for future studies on the clinical relevance and distribution of other genetic elements.

## Introduction

*Escherichia coli* is one of the most known and studied organisms, serving as a valuable bacterial model species and laboratory workhorse for molecular biology. While most *E. coli* laboratory strains do not possess pathogenic properties, *E. coli* also exhibits an extreme intra-species diversity due to its rich and variable accessory genome (1,2). This leads to some *E. coli* strains occurring as gut commensals (3,4), while others act as (opportunistic) pathogens that can cause either intestinal infections or extra-intestinal infections (in both humans and animals). Intestinal pathogenic *E. coli* (IPEC) infections display a diverse range of clinical manifestations, from mild diarrhea to potentially deadly hemorrhagic uremic syndrome (5,6). Extraintestinal pathogenic *Escherichia coli* (ExPEC) are responsible for the majority of urinary tract infections (UTIs) in humans and a large part of blood-stream infections (BSIs), which can result in sepsis (7,8). Within these two groups, different pathogenic variants or pathovars have been identified (2,9). The IPEC pathogroup comprises enteropathogenic (EPEC), shiga toxin-producing (STEC), enterohemorrhagic (EHEC), enteroaggregative (EAEC), enterotoxigenic (ETEC), and Enteroinvasive (EIEC) *E. coli*, with typical virulence factors used as specific genetic markers for these pathovars (2,10). ExPEC contain uropathogenic (UPEC), neonatal meningitis (NMEC), sepsis-associated (SEPEC), and avian pathogenic (APEC) *E. coli*. In contrast to IPEC, this subdivision is based on the clinical manifestation and source of isolation of strains, but generally without pathotype-deterministic virulence factors (2). In addition, there are specific instances in which no specific label for the associated ExPEC pathovars exists (e.g., in the case of skin and soft tissue infections or pneumonia). While ExPEC and IPEC pathogroups are typically mutually exclusive, exceptions occur (2). For example, diffuse adhering *E. coli* (DAEC) is associated with persistent diarrhea in children with Crohńs disease but also causes urinary tract infections (2). Moreover, multiple strains with hybrid pathovars have been described, including combinations of IPEC and ExPEC pathovars. Examples of this are the hybrid EAEC/STEC strain responsible for a deadly outbreak in 2011 in Germany (11) or the EAEC/UPEC hybrid detected in Australian patients with a UTI (12).

To account for the large intra-species diversity, several typing schemes have been developed for *E. coli*. One broadly used scheme divides *E. coli* into phylogenetic groups (PGs) using 17 housekeeping genes (13). In a clinical context, the Achtman multilocus sequence typing scheme, based on the allelic variants of seven house-keeping genes (14), divides *E. coli* strains into sequence types (STs) and has proven to track the typical ExPEC lineages effectively (15). In total, the Achtman scheme currently accounts for more than 10000 STs, but the majority of ExPEC infections (>85%) are linked to only 20 STs (15), with ST131 being the most represented. While several STs can also be linked to an IPEC pathotype, with ST11 being the most notable one, a serotyping scheme that considers variants of surface antigens (O for lipopolysaccharide and H for flagella) is more broadly used in the IPEC field. The most notorious IPEC serotype is O157:H7 (2,16,17), which in the majority of cases corresponds to ST11. While IPEC and their subgroups can be successfully identified and classified based on the profile of their VAGs (2,10), multiple attempts to define a similar scheme for ExPEC have proven unreliable (2,15).

*E. coli* infections have an enormous impact on human and animal health, our healthcare system, and our economy (2,6,15–18). For example, 73000 yearly cases of intestinal illnesses linked solely to *E. coli* O157:H7 were reported in US in between 1982 and 2002, with 350 individual outbreaks and more than 17% of cases requiring hospitalization (17). For ExPEC, the yearly incidence of infections in the US was around 9 million in 2003, with direct associated costs reaching 2 billion USD (19). As ExPEC lineages have only increased globally since then (15), the current impact is likely even larger. Furthermore, ExPEC are also prominent opportunistic pathogens of other mammals and birds (20), causing significant issues in the animal food-production sector and concerns regarding the One Health approach (6,21,22). Similarly, food-producing animals suffer from IPEC-related diseases and livestock can serve as a reservoir for IPEC strains, with 75% of O157:H7 human outbreaks being traced back to cattle (6,23). In addition, *E. coli* and especially ExPEC are commonly linked with antibiotic resistance genes (ARGs), which may greatly contribute to treatment failures in human and veterinary medicine and enhance the spread of antimicrobial resistance (AMR) (2,6). Despite the significant burden that ExPEC and IPEC impose on our society, and despite ExPEC being the number one cause of healthcare-associated infections, bloodstream infections and sepsis in the US (8,24–26), the pathogenic strains of *E. coli* have often been somewhat neglected and have not always received adequate attention (2). This is reflected in tendencies to leave out *E. coli*, at least initially, from important bacterial surveillance schemes such as those of the US Centers of Disease Control (CDC) or the so-called ESKAPE pathogens (2).

At the moment, it remains unclear what determines pathogenicity and pathogenic efficiency across different *E. coli* strains. For example, it is not well understood how a small proportion of *E. coli* STs, including the notorious ST131 (7,15), became responsible for the majority of global ExPEC infections. The answers to such questions may lie in large genome collections, such as EnteroBase (EB), which is the most comprehensive MLST-based *E. coli* genome collection (27). Previous studies have already demonstrated the advantages of such genomic analyses. For instance, a collection of more than 10000 *E. coli* genome sequences has been used to update the division into PGs and identify several new ones (28). Other examples include the investigation of the role of the ColV plasmid in the evolution of ST58, which belongs to the typical ExPEC STs, using 34364 draft *E. coli* genome assemblies from EB (29), and a characterization of *E. coli* pangenome diversity within and across STs using a curated dataset of 20577 *E. coli* assemblies from EB (4). However, a main limitation of these genome collections remains a lack of sufficient metadata accompanying the sequences, especially in the context of infections. This currently restricts the usability of these large collections in developing an understanding of what makes a strain pathogenic and a pathogen successful.

One factor that could contribute to pathogenicity is the presence of a Type VI secretion system (T6SS). The T6SS is a nanomachine that injects effectors into target cells, and belongs to the broader family of contractile injection systems. Effector molecules are highly diverse and target cells can be either bacteria or eukaryotic cells (30–33). As such, the contribution of T6SSs to pathogenicity can be both indirect (by mediating inter-bacterial competition to ensure niche colonization and facilitate nutrient uptake) and direct (by mediating pathogen-host interactions) (34–39). T6SSs are versatile multiprotein complexes, requiring more than 10 core proteins (32), and can be divided into four main classes: T6SS^i^, T6SS^ii^, T6SS^iii^, and T6SS^iv^. Among these, T6SS^i^ stands out as the most prevalent and it is further divided into six subclasses T6SS^i1^, T6SS^i2^, T6SS^i3^, T6SS^i4a^, T6SS^i4b^, and T6SS^i5^. Although T6SSs can be found across a wide range of Gram-negative species, including well-known pathogens from the *Pseudomonas*, *Salmonella*, *Campylobacter*, *Vibrio*, *Burkholderia*, *Serratia*, *Edwardsiella*, and *Enterobacter* genera (40–43), not all bacteria have such a secretion apparatus. In fact, differences in prevalence have been reported within the same species, and even among closely related strains (32). In the case of E. coli, three T6SS^i^ subclasses of distinct phylogenetic origin have been detected in *E. coli*: T6SS^i1^, T6SS^i2^, and T6SS^i4b^. In the *E. coli* field, they are commonly referred to as T6SS1 (corresponding to T6SS^i2^), T6SS2 (T6SS^i1^) and T6SS3 (T6SS^i4b^). In the literature, the associated nomenclature is not unified and various terms have been used to describe T6SSs and their subdivision. Terms such as types (44), forms (45), subtypes (44), class (30), clusters (44), phylogenetic groups (46), groups (47), locus (47–49), subclass (50), subgroups (51) are used apparently interchangeably. To ensure consistency within this manuscript, we stick to the terms class (T6SS^i^), subclass (T6SS^i1^, T6SS^i2^, T6SS^i4b^) and groups (referring to further subdivision of subclasses). In *E. coli*, a total of 13 core genes (*tssA*-*tssM*) are required for the T6SS^i^ functionality. These genes can be accompanied by additional genes determining the specificity of the spike (38,46), by variable (toxic) effectors, and by immunity genes that neutralize effector toxicity for cognate T6SS-harboring cells (32,46).

In this study, we aimed to assess the prevalence and distribution of different T6SSs in *E. coli*, in order to develop better insight into its potential contribution to pathogenicity in this species. While previous studies have already focused on T6SSs distribution across different PGs of *E. coli*, they represent smaller-scale studies with limited numbers of strains or strains of limited origin (e.g., considering only a subset of STs) (47,49,52,53). To develop a more comprehensive picture, we determined the prevalence and distribution of T6SS subclasses in a large ST-unbiased collection of 131610 *E. coli* genomes recovered from EnteroBase, with updated annotations regarding their clinical properties. For this, we constructed a comprehensive database of *E. coli*-relevant T6SS^i^ regions that combines input from different sources, examined their prevalence across PGs, pathogroups and pathovars, and determined their co-occurrence with other VAGs and MDR. The results of this large-scale analysis are described below.

## Results

### Composition and annotation of a large-scale *E. coli* genome collection

To uncover T6SS distribution across *E. coli*, we first put together a large-scale *E. coli* genome collection. For this, we downloaded a total of 136051 available *E. coli* genomes with metadata from EB. We focused on sequences with sufficient metadata to determine their potential pathogenic properties, and performed additional annotations to extract more clinically relevant information of the strains (see Methods). We identified PGs using the ClermonTyping tool (54), merged these results with those already present in the EB metadata, and filtered out genomes that showed irregular PG results (see Methods). After additional filtering (see Methods), we ended up with a collection of 131610 genomes. The distribution of these genomes across PGs is shown in Fig. 1A. We found that PG B1 was the most prevalent (40059 genomes) and clade I the least prevalent (489 genomes) (Fig. 1A, Table 1). Relative distributions within PGs based on the geographical location (*Continent*) of genomes indicated representation from all continents (except Antarctica), with a roughly similar distribution across PGs and a clear bias towards genomes from Europe and North America (Fig. 1B). A subdivision based on the niche from which strains were isolated (*Niche*) revealed a general bias in our collection towards genomes coming from humans, although we could still identify notable differences between PGs (Fig. 1B). For example, B2 contains a higher fraction of genomes of human origin than other PGs, but a lower fraction of livestock origin. On the other hand, PG G is enriched with genomes coming from poultry.

**Figure 1.**
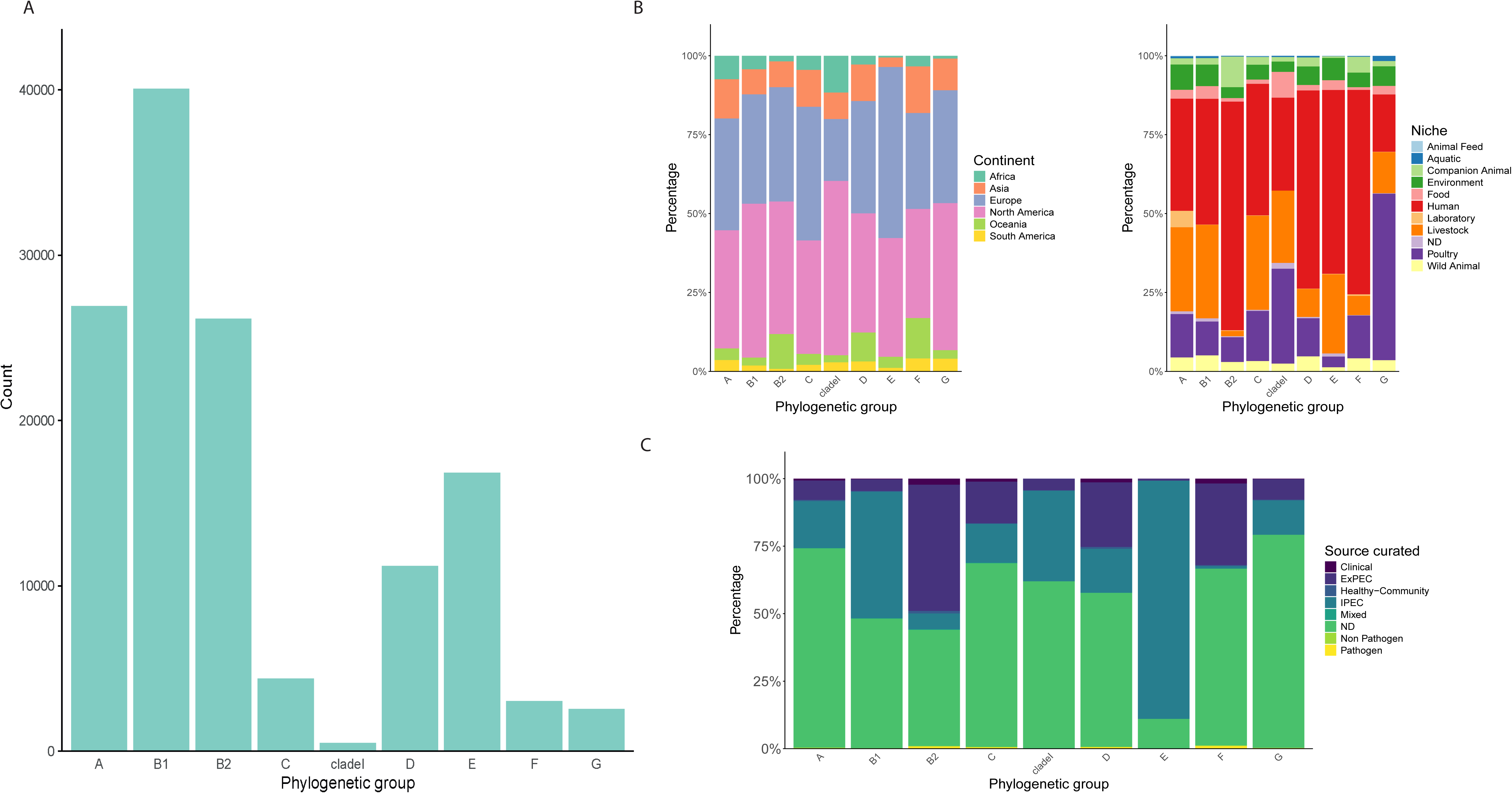
Composition of a large-scale *E. coli* genome collection. **A.** Bar graph showing the distribution of our collection of genomes (131610) across major phylogenetic groups. **B.** Stacked bar graphs showing the relative distribution of selected metadata categories, *Continent* (left) and *Niche* (right), within the phylogenetic groups. **C.** Stacked bar graph showing the relative distribution of metadata category *Source curated,* which reflects the clinical characteristics of genomes.

**Table 1.** Prevalence of T6SS^i^ subclasses (*Presence* and *Completeness*) across selected metadata categories. PT6SS^i1-i4b^ corresponds to the *Presence* of specific T6SS^i^ subclasses while CT6SS^i1-i4b^ corresponds to *Completeness*. P/C-Mul corresponds to genomes with multiple T6SS^i^ subclasses. N specifies the number of genomes in the respective categories. The same type of prevalence analysis for both DBs separately is presented in Table S3.

| ***E. coli*** | **N** | **PT6SSi1** | **PT6SSi2** | **PT6SSi4b** | **P-Mul** | **CT6SSi1** | **CT6SSi2** | **CT6SSi4b** | **C-Mul** |
| --- | --- | --- | --- | --- | --- | --- | --- | --- | --- |
|  | 131610 | 55,9% | 28,3% | 1,3% | 14,7% | 40,4% | 20,7% | 1,3% | 7,8% |
| **PG** | **N** | **PT6SSi1** | **PT6SSi2** | **PT6SSi4b** | **P-Mul** | **CT6SSi1** | **CT6SSi2** | **CT6SSi4b** | **C-Mul** |
| A | 26922 | 26,0% | 4,8% | 2,4% | 3,0% | 21,7% | 3,5% | 2,4% | 2,3% |
| B1 | 40059 | 69,5% | 22,6% | 1,9% | 20,4% | 53,0% | 19,3% | 1,8% | 15,1% |
| B2 | 26148 | 38,1% | 90,7% | 0,0% | 36,7% | 11,7% | 60,3% | 0,0% | 11,2% |
| C | 4381 | 40,6% | 3,7% | 0,0% | 3,4% | 38,6% | 3,1% | 0,0% | 2,9% |
| cladel | 489 | 44,2% | 3,1% | 0,0% | 2,0% | 22,9% | 3,1% | 0,0% | 1,8% |
| D | 11207 | 81,7% | 2,5% | 2,1% | 2,5% | 64,9% | 2,2% | 2,1% | 2,1% |
| E | 16829 | 95,6% | 0,8% | 0,3% | 0,8% | 74,8% | 0,7% | 0,3% | 0,4% |
| F | 3031 | 42,7% | 0,3% | 0,0% | 0,1% | 39,2% | 0,1% | 0,0% | 0,0% |
| G | 2544 | 9,1% | 98,5% | 0,0% | 8,4% | 8,6% | 90,4% | 0,0% | 7,7% |
| **Niche** | **N** | **PT6SSi1** | **PT6SSi2** | **PT6SSi4b** | **P-Mul** | **CT6SSi1** | **CT6SSi2** | **CT6SSi4b** | **C-Mul** |
| Animal Feed | 145 | 47,6% | 9,0% | 0,0% | 5,5% | 38,6% | 6,9% | 0,0% | 4,1% |
| Aquatic | 604 | 48,3% | 24,7% | 0,2% | 13,7% | 32,3% | 22,7% | 0,2% | 4,8% |
| Companion Animal | 4595 | 61,8% | 58,3% | 0,1% | 36,5% | 29,3% | 53,7% | 0,1% | 11,8% |
| Environment | 8171 | 55,0% | 18,2% | 0,6% | 9,5% | 41,6% | 14,6% | 0,6% | 5,0% |
| Food | 3569 | 62,1% | 18,5% | 0,1% | 9,2% | 48,3% | 13,0% | 0,1% | 6,2% |
| Human | 65700 | 56,4% | 35,7% | 2,4% | 17,2% | 39,1% | 23,2% | 2,4% | 9,4% |
| Laboratory | 1566 | 6,2% | 5,1% | 0,0% | 2,0% | 4,7% | 4,6% | 0,0% | 0,9% |
| Livestock | 26615 | 59,9% | 12,9% | 0,0% | 9,3% | 51,4% | 11,0% | 0,0% | 6,8% |
| ND | 965 | 56,2% | 16,6% | 0,6% | 12,4% | 38,7% | 14,8% | 0,6% | 7,7% |
| Poultry | 14571 | 47,8% | 27,0% | 0,0% | 12,8% | 30,6% | 25,1% | 0,0% | 4,3% |
| Wild Animal | 5109 | 58,9% | 21,7% | 0,2% | 13,1% | 43,0% | 18,9% | 0,2% | 6,3% |
| **Source curated** | **N** | **PT6SSi1** | **PT6SSi2** | **PT6SSi4b** | **P-Mul** | **CT6SSi1** | **CT6SSi2** | **CT6SSi4b** | **C-Mul** |
| Clinical | 1136 | 46,0% | 54,6% | 3,0% | 19,0% | 29,7% | 29,4% | 3,0% | 8,1% |
| ExPEC | 20523 | 50,0% | 60,4% | 0,3% | 24,5% | 27,4% | 40,1% | 0,3% | 8,4% |
| Healthy-Communit | 571 | 49,6% | 46,1% | 0,2% | 20,5% | 33,6% | 38,0% | 0,2% | 9,5% |
| IPEC | 42791 | 71,0% | 15,4% | 0,1% | 13,1% | 57,5% | 13,3% | 0,1% | 10,2% |
| Mixed | 159 | 34,6% | 6,9% | 0,0% | 2,5% | 24,5% | 4,4% | 0,0% | 2,5% |
| ND | 65828 | 48,2% | 25,9% | 2,3% | 12,6% | 33,8% | 19,3% | 2,3% | 6,0% |
| Non Pathogen | 183 | 41,5% | 27,3% | 0,5% | 7,1% | 33,3% | 22,4% | 0,5% | 2,2% |
| Pathogen | 419 | 45,8% | 52,5% | 5,0% | 15,8% | 30,5% | 19,6% | 5,0% | 6,9% |
| **ExPEC type** | **N** | **PT6SSi1** | **PT6SSi2** | **PT6SSi4b** | **P-Mul** | **CT6SSi1** | **CT6SSi2** | **CT6SSi4b** | **C-Mul** |
| APEC | 446 | 50,4% | 46,6% | 0,2% | 20,6% | 28,0% | 44,8% | 0,2% | 6,5% |
| BSI | 4957 | 40,9% | 66,4% | 0,3% | 18,7% | 23,6% | 33,6% | 0,3% | 5,8% |
| ExPEC | 2524 | 51,1% | 40,9% | 0,0% | 18,6% | 31,5% | 28,4% | 0,0% | 7,0% |
| No | 42608 | 71,1% | 15,4% | 0,1% | 13,1% | 57,6% | 13,3% | 0,1% | 10,3% |
| NS | 68322 | 48,2% | 26,7% | 2,3% | 12,8% | 33,7% | 19,6% | 2,3% | 6,0% |
| Respiratory | 407 | 41,5% | 64,1% | 0,0% | 20,4% | 23,1% | 33,2% | 0,0% | 7,6% |
| UPEC | 12346 | 53,6% | 61,6% | 0,3% | 28,0% | 28,2% | 44,7% | 0,3% | 9,8% |
| **IPEC type** | **N** | **PT6SSi1** | **PT6SSi2** | **PT6SSi4b** | **P-Mul** | **CT6SSi1** | **CT6SSi2** | **CT6SSi4b** | **C-Mul** |
| EHEC | 24298 | 76,3% | 8,3% | 0,0% | 8,3% | 63,7% | 8,2% | 0,0% | 7,9% |
| EIEC | 356 | 76,4% | 1,7% | 0,0% | 0,6% | 49,4% | 1,1% | 0,0% | 0,0% |
| EPEC | 5176 | 50,9% | 8,7% | 0,0% | 8,5% | 36,1% | 8,5% | 0,0% | 7,7% |
| ETEC | 2266 | 51,1% | 3,8% | 0,0% | 3,1% | 42,6% | 3,4% | 0,0% | 1,9% |
| ETEC/EHEC | 102 | 9,8% | 0,0% | 0,0% | 0,0% | 6,9% | 0,0% | 0,0% | 0,0% |
| ETEC/EPEC | 46 | 26,1% | 0,0% | 0,0% | 0,0% | 0,0% | 0,0% | 0,0% | 0,0% |
| ETEC/STEC | 557 | 38,1% | 1,3% | 0,0% | 1,1% | 18,1% | 1,3% | 0,0% | 0,4% |
| ND | 501 | 59,1% | 31,3% | 0,4% | 19,8% | 31,7% | 27,7% | 0,4% | 7,0% |
| No | 88342 | 48,6% | 34,5% | 1,8% | 15,5% | 32,2% | 24,3% | 1,8% | 6,6% |
| STEC | 9966 | 75,3% | 40,1% | 0,6% | 30,8% | 59,9% | 31,6% | 0,5% | 20,3% |

We also subdivided the genomes based on their clinical relevance (*Source curated*). This subdivision consisted of three main groups: not determined (*ND*), *IPEC*, and *ExPEC* (Fig. 1C). The largest one, *ND*, comprised about half of our collection (65828 genomes) and represents genomes for which this additional annotation could not be assigned based on the available metadata. While this group is expected to contain less pathogenic strains than the other two, it cannot be perceived as synonymous with non-pathogenic, and likely houses representatives of all possible lifestyles of *E. coli*, from environmental strains to commensals and pathogens (of unknown proportions/severity). The *IPEC* group consisted of 42791 genomes, mostly assigned using the VAGs-based scheme (97% of IPEC genomes). We assigned genomes to the *ExPEC* group (20523 genomes) mainly based on the source from which they were isolated. We hereby focused on the presence of *E. coli* in areas associated with symptomatic or asymptomatic infections (e.g., blood or urine). Other subgroups related to the clinical relevance of genomes were smaller, ranging from 1136 genomes identified as *Clinical* (referring to an unspecific clinical annotation) to 159 genomes identified as *Mixed* (referring to mixed IPEC-ExPEC pathotype). The other minor groups were represented by *Healthy-Community* (571 genomes), *Pathogen* (419 genomes), and *Non Pathogen* (183 genomes). Examination of this subdivision across PGs revealed an increased fraction of IPEC genomes in PG B1 and E, while an increased fraction of ExPEC genomes was observed in PG B2, F and D (Fig. 1C).

The *ExPEC* and *IPEC* group were also divided into several subgroups (see Methods) and further investigated separately (Tables S4 and S5). This analysis revealed an uneven distribution of the subgroups across PGs, especially apparent for IPEC (Fig. S2). *EHEC* represents the major pathovar of PG E with a small fraction of other pathovars in this PG, while PG B1 contained similar proportions of *EHEC* and *STEC*, and a considerable fraction of *EPEC* (Fig. S2). We also examined which STs are linked with *ExPEC* (Table S6) or *IPEC* (Table S7), and assessed their distribution across *ExPEC* (Table S4) and *IPEC* (Table S5) types. This analysis confirmed the role of ST131 as the major ExPEC ST with 24% of ExPEC-labelled strains corresponding to this ST and only *APEC* not having ST131 as the most common ST (ST117 was the most dominant ST within this *ExPEC* type). Besides ST131, only one other ST represented more than 5% of *ExPEC* strains, namely ST73 with 7%. Within IPEC, the dominance of ST11 is apparent as it represents almost one third (33%) of IPEC genomes. The only other *IPEC* ST reaching more than 5% is ST21 (9%). Across *IPEC* pathovars, ST11 mostly dominated *EHEC* and was the second most prevalent ST for *EPEC*, after ST10.

### Construction of a comprehensive database of *E. coli* T6SS components reveals the distribution of T6SS^i^ subclasses across *E. coli* genomes

To investigate the distribution of various T6SS^i^ subclasses across our *E. coli* genome collection, we aimed to construct a comprehensive sequence database of *E. coli* T6SS components. For this, we combined sequences from different available sources (see Methods), and focused on genes annotated as one of the 13 core genes (*tssA*-*tssM*). Using this database, we evaluated the presence of these genes for each T6SS^i^ subclass across our *E. coli* genome collection. The frequency distributions of the number of detected *tss* core genes revealed an almost binary pattern for each T6SS^i^ subclass (Fig. 2A), where either (almost) all components of a given T6SS^i^ subclass were present, or none. We exploited this distribution pattern to assess both the completeness and presence of the three T6SS^i^ subclasses that occur in *E. coli*. *Present/Presence* (P) corresponds to the detection of at least 12 core genes from the respective subclass, and *Complete/Completeness* (C) to the detection of all 13 core genes, where we expect the corresponding T6SS to be functional. We chose to include both, as functionality might still be preserved for some members in the *Present* group (given that our database, although comprehensive, will never be perfectly complete and some of the core genes components can be found in orphan regions (40,46)).

**Figure 2.**
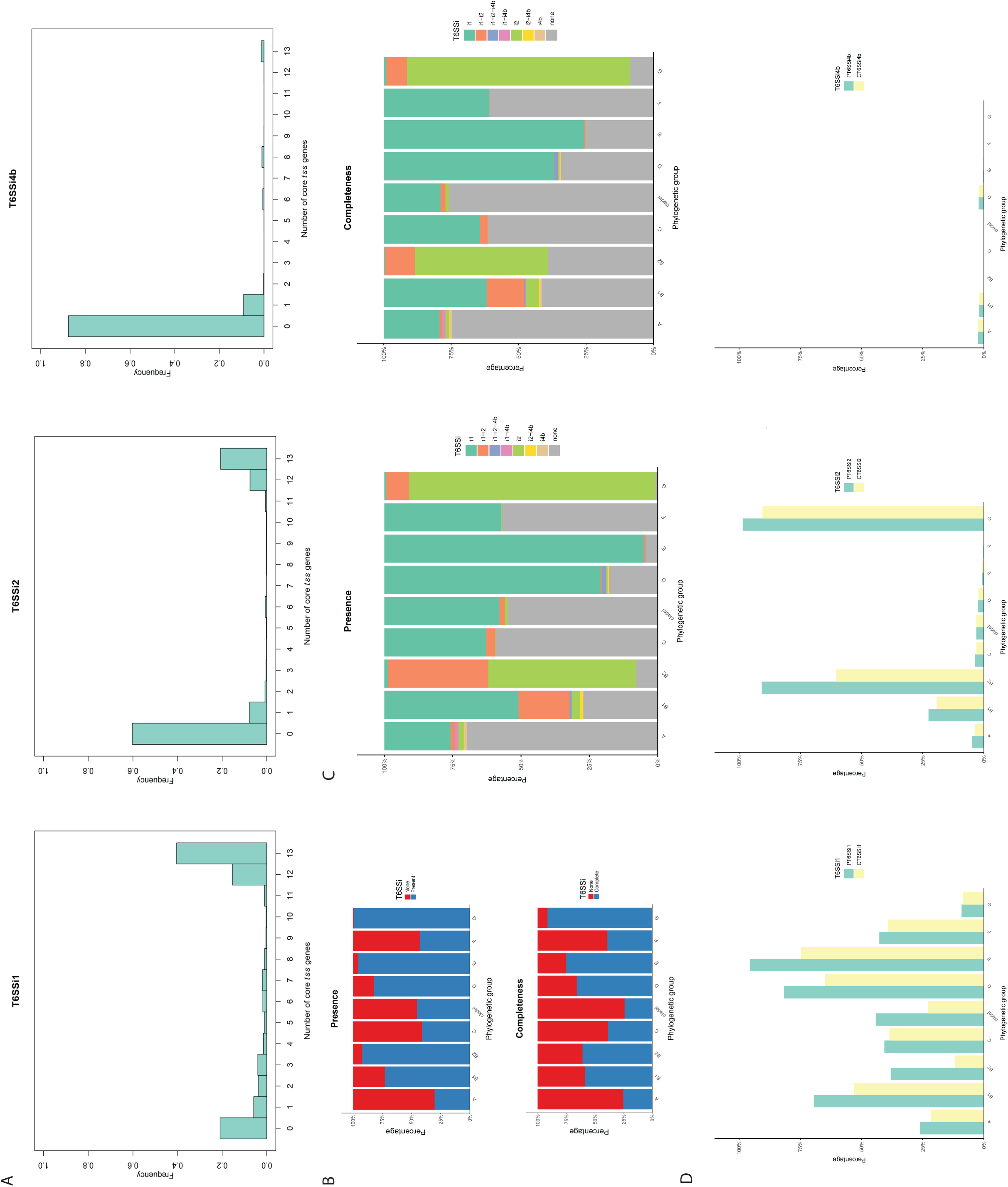
T6SS^i^ subclasses in the collection of genomes. **A.** Bar graphs showing the fraction of genomes with the indicated number of core *tss* genes (0-13) for T6SS^i1^ (left), T6SS^i2^ (middle) and T6SS^i4^ (right). **B.** Stacked bar graphs showing the percentage of genomes with at least one subclass of T6SS^i^ detected as *Present* (top) or *Complete* (bottom) across the phylogenetic groups. **C.** Stacked bar graphs showing the specific combinations of T6SS^i^ subclasses that were detected as *Present* (left) or *Complete* (right) across the phylogenetic groups. **D.** Bar graphs showing the *Presence* (green bars) and *Completeness* (yellow bars) of T6SS^i1^ (left), T6SS^i2^ (middle) and T6SS^i4^ (right) across the phylogenetic groups.

Upon examination of the *Presence* and *Completeness* of T6SSs across *E. coli* genomes, we found that more than half of the genomes contained at least one T6SS (70.3% and 54.3% for P and C, respectively; Fig. 2B). Moreover, we identified a substantial fraction of genomes containing multiple T6SS^i^ subclasses with 14.7% and 7.8% for P and C, respectively (Fig. 2C). Most of the genomes with multiple T6SS^i^ subclasses combined T6SS^i1^ and T6SS^i2^ (14.0% and 7.2% for P and C), but other combinations also occurred (Fig. 2C). A more detailed investigation of the distribution across PGs revealed distinct distribution patterns between subclasses (Fig. 2D). While T6SS^i1^ was broadly distributed, and was prevalent across all PGs, the distribution of both T6SS^i2^ and T6SS^i4b^ appeared more limited, to only a subset of PGs. T6SS^i2^ was found in a relatively large fraction of genomes from PGs B1, B2, and G, while T6SS^i4b^ was not very prevalent and only found in a small fraction of genomes of PGs A, B1, and D. In general, we found a good agreement between the *Presence* of and *Completeness* of the T6SS subclasses across PGs, with the largest differences observed in PG B2 for both T6SS^i1^ and T6SS^i2^. As a control, we also performed the same analysis using the separate sequences of T6SS components, based on the database they originated from. This analysis revealed the added value of our merged database, as sequences retrieved from separate databases often only partially recapitulated the global distribution pattern of different T6SS^i^ subclasses (Fig. S3). For example, the T6SS^i1^ was almost exclusively linked to B2 in one of the databases (Fig. S3), which is likely an artefact of focusing only on a subset of sequences, given that this association no longer holds when considering all T6SS^i1^ sequences (Fig. 2D).

To gain further insight into the potential association of T6SSs with pathogenic properties of strains, we also investigated their distribution across the *Source curated* category of genomes. This analysis revealed an enrichment of T6SS^i1^ for *IPEC* genomes (Fig. 3A and Table S8). For T6SS^i2^, the *ExPEC* genomes were the largest group, although this proportion was statistically indistinguishable from the *Healthy-Community* group for the *Complete* T6SS^i2^ (p-value < 0.31, Table S8). Within the T6SS^i2^-positive genomes, we observed a strong depletion of *IPEC* and *Mixed* genomes in comparison to other groups (Fig. 3A and Table S8). Differences among groups could also be observed for T6SSi^4b^ (Fig. 3A), yet the low absolute values do not allow T6SSi^4b^ to be a relevant marker for any clinical group. In a subsequent step, we explored associations of T6SS^i^ subclasses with *ExPEC* and *IPEC* types (Fig. 3B). All *ExPEC* groups displayed an enrichment in T6SS^i2^ compared to other groups (Table S8), with *APEC* and *UPEC* showing the highest association (Fig. 3B). In contrast, for the *IPEC* group, all major pathovars displayed an enrichment in T6SS^i1^ with the comparative group (*No*; Table S8), with the exception of the minor hybrid groups (*ETEC/EHEC*, *ETEC/EPEC*, *ETEC/STEC*). *EHEC* and *STEC* contained the largest fraction of genomes with T6SS^i1^ (Fig. 3C), while *STEC* showed an, for IPEC, atypical association with T6SS^i2^ (Fig. 3C and Table S8).

**Figure 3.**
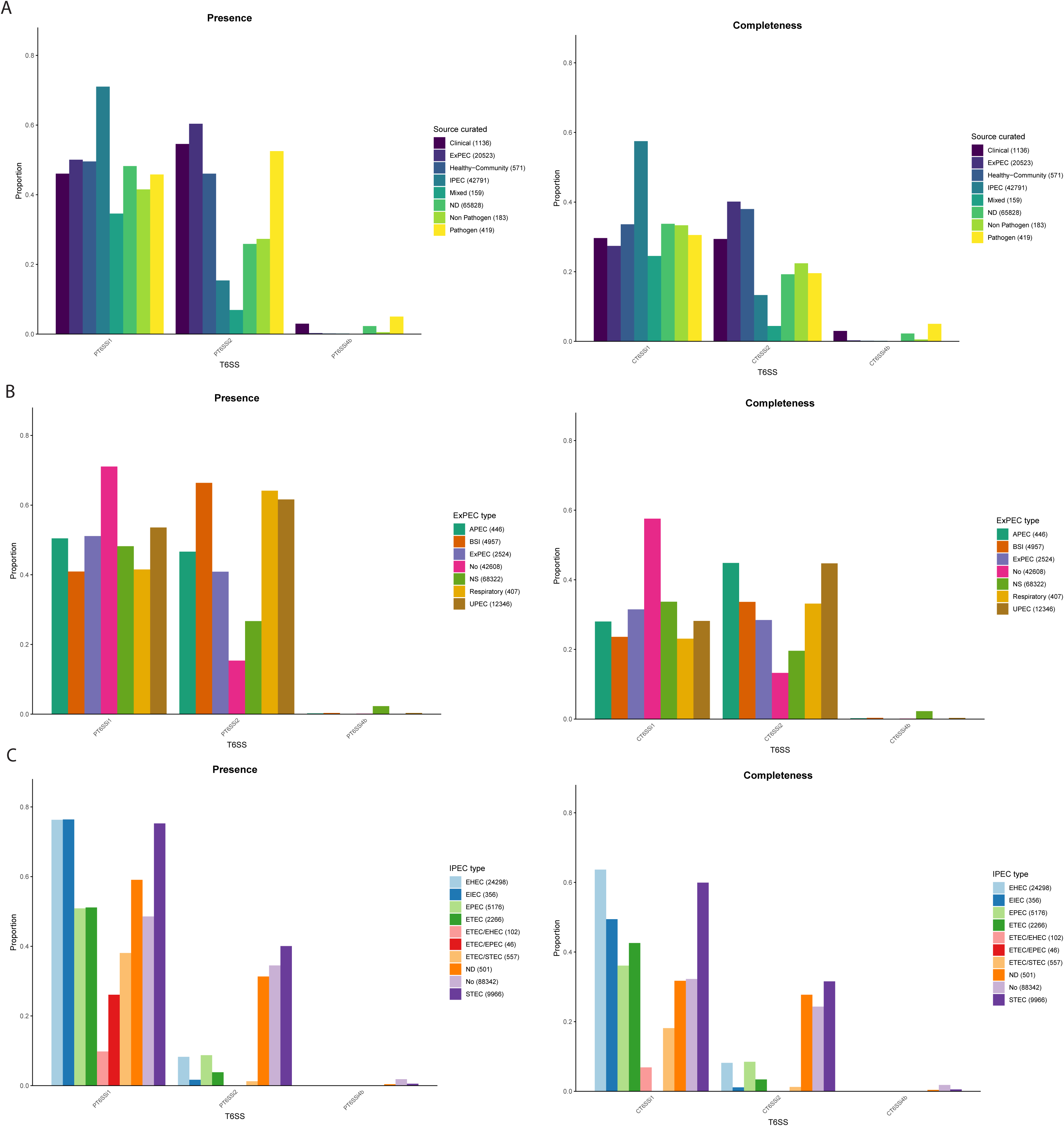
Proportions of T6SSi subclasses in clinically relevant groups. For all panels, the proportion of T6SS^i^ subclasses that were detected as *Present* are shown on the left, while those that were detected as *Complete* are shown on the right. **A.** Bar graphs showing the proportions of T6SS^i^ subclasses across the *Source curated* category. **B.** Bar graphs showing the proportions of T6SS^i^ subclasses across ExPEC types. **C.** Bar graphs showing the proportions of T6SS^i^ subclasses across IPEC types.

### Distribution of T6SS^i^ subclasses across major *E. coli* STs

In a subsequent step, we examined the distribution of T6SS^i^ subclasses across the 35 most prevalent STs of *E. coli* (each containing >500 genomes). An overview of this analysis is presented in Fig. 4. In general, T6SS^i^ subclasses displayed a more delineated distribution pattern, with stronger enrichments and absences within STs than within PGs. In the majority of cases, a given T6SS^i^ subclass was either present or not within an ST (Fig. 4 and Table 2). With the exception of ST155, T6SS^i1^ is either *Present* (>85%) or almost completely absent (<20%) for specific STs. This absence is not linked to specific PGs, as T6SS^i1^ is not present in ST131 and ST73 (B2), ST32 (D), ST744 (A) or ST21 (B1). ST21 is particularly relevant, given that it represents the second most dominant *IPEC* ST, yet shows no presence of any T6SS^i^. T6SS^i2^ was present in all STs in PG B2 (ST131, ST1193, ST127, ST73, ST372, ST95 and ST12) and the only ST from G (ST117), which all belong to prominent ExPEC. At the same time, T6SS^i2^ was also prevalent in two STs from the B1 PG linked to an IPEC pathotype, namely ST17 and ST442. We only found T6SS^i4b^ in a fraction of ST34 genomes, where they typically co-occurred with T6SS^i2^. Strains belonging to several other STs, including ST12, ST1193, ST17, and ST442, also contained multiple T6SSs, which typically combined T6SS^i1^ and T6SS^i2^ (Fig. 4).

**Figure 4.**
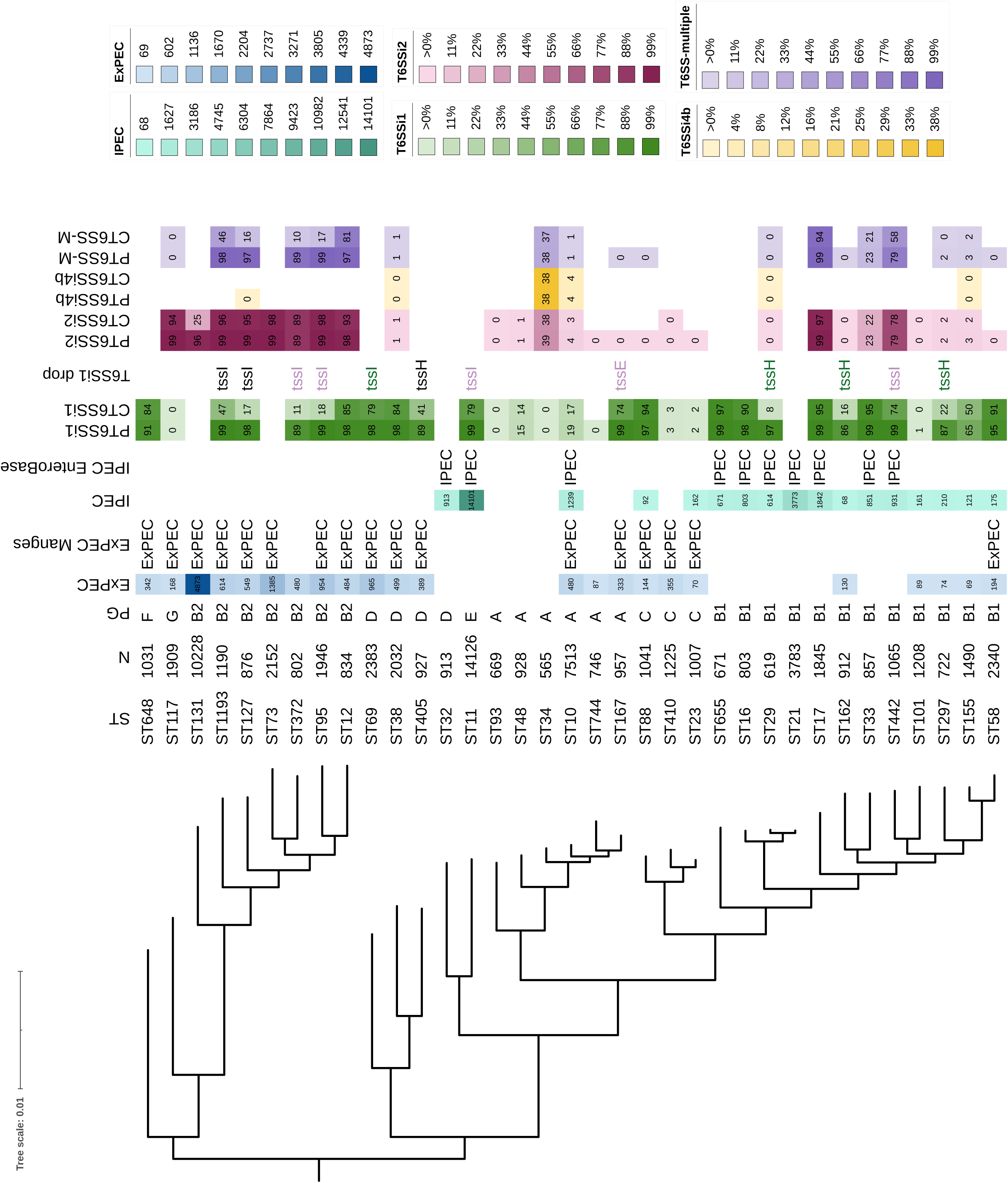
Phylogenetic analysis of 35 most dominant STs. The metadata included in our phylogenetic tree are sequence types (**ST**), number of genomes of the respective ST in our collection (**N**), phylogenetic group (**PG**), heatmap highlighting the dominant ExPEC STs in our study (threshold of min. 50 ExPEC genomes) **(EXPEC)**, ExPEC label given to 20 dominant ExPEC STs undertaken from(15) (**ExPEC Manges**), heatmap highlighting the dominant IPEC STs (threshold of min. 50 IPEC genomes) (**IPEC**), IPEC label given to STs which belonged among the 20 dominant IPEC STs determined based on EnteroBase information (**IPEC EnteroBase**), heatmap showing the prevalence (%) of T6SS^i1^ for each ST (**PT6SSi1, CT6SSi1**), T6SS^i1^-associated *tss* genes that could be linked to the drop between P and C (**T6SSi1 drop**), – the genes in black were not found, the genes in purple were found truncated and the genes in green were identified after additional examination of the T6SS contigs heatmap showing the prevalence of T6SS^i2^ for each ST (**PT6SSi2, CT6SSi2**), heatmap showing the prevalence of T6SS^i4b^ for each ST (**PT6SSi4b, CT6SSi4b**), heatmap showing the prevalence of multiple T6SS^i^ subclasses for each ST (**PT6SS-M, CT6SS-M**). P/C corresponds to *Present* and *Complete*, respectively.

**Table 2.** Prevalence of T6SS^i^ subclasses (*Presence* and *Completeness*) in the 35 dominant STs. The 35 STs were selected from our dataset based on being represented by more than 500 genomes. PT6SS^i1-i4b^ correspond to the *Presence* of specific T6SS^i^ subclass, while CT6SS^i1-i4b^ correspond to *Completeness*, P/C-Mul considers genomes with multiple T6SS^i^ subclasses, and N specifies the number of genomes in the respective category.

| ST | N | PT6SSi1 | PT6SSi2 | PT6SSi4b | P-Mul | CT6SSi1 | CT6SSi2 | CT6SSi4b | C-Mul |
| --- | --- | --- | --- | --- | --- | --- | --- | --- | --- |
| ST11 | 14126 | 99,4% | 0,0% | 0,0% | 0,0% | 79,5% | 0,0% | 0,0% | 0,0% |
| ST131 | 10228 | 0,0% | 96,3% | 0,0% | 0,0% | 0,0% | 25,0% | 0,0% | 0,0% |
| ST10 | 7513 | 19,9% | 4,8% | 4,0% | 1,8% | 17,9% | 3,2% | 4,0% | 1,6% |
| ST21 | 3783 | 0,0% | 0,0% | 0,0% | 0,0% | 0,0% | 0,0% | 0,0% | 0,0% |
| ST69 | 2383 | 98,9% | 0,0% | 0,0% | 0,0% | 79,9% | 0,0% | 0,0% | 0,0% |
| ST58 | 2340 | 95,3% | 0,6% | 0,0% | 0,6% | 91,7% | 0,0% | 0,0% | 0,0% |
| ST73 | 2152 | 0,0% | 99,3% | 0,0% | 0,0% | 0,0% | 98,3% | 0,0% | 0,0% |
| ST38 | 2032 | 98,7% | 1,7% | 0,6% | 1,7% | 84,9% | 1,3% | 0,6% | 1,3% |
| ST95 | 1946 | 99,5% | 99,8% | 0,0% | 99,4% | 18,0% | 98,7% | 0,0% | 17,8% |
| ST117 | 1909 | 0,1% | 99,6% | 0,0% | 0,1% | 0,1% | 94,0% | 0,0% | 0,1% |
| ST17 | 1845 | 99,3% | 99,2% | 0,0% | 99,1% | 95,6% | 97,9% | 0,0% | 94,5% |
| ST155 | 1490 | 65,7% | 3,0% | 0,3% | 3,2% | 50,1% | 2,3% | 0,3% | 2,6% |
| ST410 | 1225 | 3,9% | 0,1% | 0,0% | 0,0% | 3,8% | 0,1% | 0,0% | 0,0% |
| ST101 | 1208 | 1,9% | 0,2% | 0,0% | 0,0% | 0,8% | 0,1% | 0,0% | 0,0% |
| ST1193 | 1190 | 99,0% | 99,3% | 0,0% | 98,8% | 47,2% | 96,1% | 0,0% | 46,6% |
| ST442 | 1065 | 99,1% | 79,5% | 0,0% | 79,2% | 74,1% | 78,2% | 0,0% | 58,0% |
| ST88 | 1041 | 97,9% | 0,9% | 0,0% | 0,9% | 94,4% | 0,0% | 0,0% | 0,0% |
| ST648 | 1031 | 91,5% | 0,0% | 0,0% | 0,0% | 84,2% | 0,0% | 0,0% | 0,0% |
| ST23 | 1007 | 2,9% | 0,4% | 0,0% | 0,0% | 2,7% | 0,0% | 0,0% | 0,0% |
| ST167 | 957 | 99,2% | 0,1% | 0,0% | 0,1% | 74,5% | 0,0% | 0,0% | 0,0% |
| ST48 | 928 | 15,2% | 1,0% | 0,0% | 0,0% | 14,4% | 1,0% | 0,0% | 0,0% |
| ST405 | 927 | 89,4% | 0,0% | 0,0% | 0,0% | 41,9% | 0,0% | 0,0% | 0,0% |
| ST32 | 913 | 0,0% | 0,0% | 0,0% | 0,0% | 0,0% | 0,0% | 0,0% | 0,0% |
| ST162 | 912 | 86,6% | 0,5% | 0,0% | 0,3% | 16,2% | 0,2% | 0,0% | 0,0% |
| ST127 | 876 | 98,1% | 99,3% | 0,1% | 97,4% | 17,1% | 95,9% | 0,0% | 16,3% |
| ST33 | 857 | 99,1% | 23,7% | 0,0% | 23,7% | 95,4% | 22,6% | 0,0% | 21,7% |
| ST12 | 834 | 98,4% | 98,1% | 0,0% | 97,0% | 85,1% | 93,9% | 0,0% | 81,2% |
| ST16 | 803 | 98,8% | 0,0% | 0,0% | 0,0% | 90,3% | 0,0% | 0,0% | 0,0% |
| ST372 | 802 | 89,7% | 89,9% | 0,0% | 89,2% | 11,0% | 89,2% | 0,0% | 10,2% |
| ST744 | 746 | 0,1% | 0,1% | 0,0% | 0,0% | 0,0% | 0,0% | 0,0% | 0,0% |
| ST297 | 722 | 87,8% | 2,8% | 0,0% | 2,6% | 22,0% | 2,2% | 0,0% | 0,7% |
| ST655 | 671 | 99,9% | 0,0% | 0,0% | 0,0% | 97,8% | 0,0% | 0,0% | 0,0% |
| ST93 | 669 | 0,1% | 0,1% | 0,0% | 0,0% | 0,1% | 0,1% | 0,0% | 0,0% |
| ST29 | 619 | 97,1% | 0,6% | 0,2% | 0,8% | 80,0% | 0,6% | 0,2% | 0,8% |
| ST34 | 565 | 0,5% | 39,5% | 38,9% | 38,8% | 0,2% | 38,8% | 38,6% | 37,9% |

As before, we found a good concordance between the *Presence* and *Completeness* of T6SS^i^ subclasses, with some notable exceptions (Fig. 4). While T6SS^i2^ is present in the majority of ST131 genomes, only 25% contain a complete T6SS^i2^. This drop has been described in a previous study in which it was linked to an interruption of the *tssM* gene, encoding part of the T6SS membrane complex, by IS*Ec12* (53), a mobile genetic element. For T6SS^i1^, we could identify several STs that displayed large discrepancies (≥ 15%) between the *Presence* and *Completeness* of this subclass. With the exception of ST155, the drop from P to C could typically be attributed to a single *tss* gene per ST (Fig. 4). Inspection of the relevant region revealed three possible scenarios: (i) the missing component was present and intact in the region (ST69, ST29, ST162 and ST297), but its sequence was not present in our T6SS component database (a likely consequence of our database only including experimentally validated T6SSs), (ii) the missing gene was not identified in the contigs carrying the *tss* genes (ST1193, ST127, ST405), suggesting it was lost, part of an orphan region not included in the T6SS databases, or a bioinformatically predicted T6SS gene located on a different contig, or (iii) the missing gene was truncated (ST372, ST95, ST11, ST167 and ST442) (Fig. 4 and S8). For example, a residue of the IS*3* family transposase was recovered next to the *tssE* of ST167, likely playing role in its truncation in this particular case.

To highlight the potential clinical relevance of the 35 dominant STs, we annotated them with an ExPEC/IPEC label and indicated the amount of genomes within the ST that we assigned to the *ExPEC* or *IPEC* category (Fig. 4). The ExPEC label was awarded based on a previous meta-analysis of more than 100 ExPEC studies to determine the most dominant ExPEC STs (15). In general, our own determinations of the number of ExPEC genomes within an ST largely corresponded to these labels, with some notable exceptions. For example, we identified elevated fractions of ExPEC strains in several STs that do not belong to the dominant 20 STs responsible for the large majority of ExPEC infections. This included ST372, but also ST744, ST162, ST101, and ST297. For IPEC, we awarded the IPEC label to STs that belonged to the 20 most dominant IPEC STs determined by our analysis based largely on the virulence scheme present in EB (27). Similar to ExPEC, the fraction of IPEC genomes within an ST largely corresponded to these labels, with ST23, ST101, ST297, ST155, and ST58 as exceptions. Only specific STs displayed elevated fractions of both ExPEC and IPEC genomes, with ST10 being the most notable example. Further analysis revealed that this observation typically stems from two distinct proportions of strains within an ST (exclusively ExPEC and exclusively IPEC), rather than from a hybrid pathotype, yet this is not conclusive as a large proportion of the genomes is labelled as *ND* (Fig. S9).

### T6SS-related traits of *E. coli* ST131 and ST11

After mapping the general distribution pattern of T6SSs in *E. coli*, we set out to investigate T6SS-related properties of important pathogenic strains in more detail. For ExPEC, we focused on ST131, the major ExPEC ST. This ST also contained a unique pattern of reduced T6SS^i2^ *Completeness* (Fig. 4), which was previously traced back to a *tssM* fragmentation (53). To examine the prevalence of this fragmentation across ST131, we evaluated the presence of ST131-specific *tssM* fragments (A, B) and IS*Ec12*. This analysis revealed that either the intact *tssM* sequence was present (likely generating a functional T6SS) or both *tssM* fragments and the IS*Ec12* were present (likely generating a non-functional T6SS) (Fig. 5A). We only found other options in very rare instances (Fig. 5A). Fragmentation of *tssM* in ST131 did display a distinct niche-related pattern (Fig. 5A). While some categories were limited in terms of representation and could not reliably be compared to others (i.e, *Aquatic* or *Laboratory*), we found a significant difference between *Humans* and *Poultry* (p-value < 10^-209^) or *Humans* and *Livestock* (p-value < 10^-61^). Whereas *tssM* was largely fragmented in *Humans*, it was mostly intact in genomes belonging to the *Poultry* or *Livestock* categories. Besides this potential *tssM* fragmentation, this region also contained three *vgrG* (*tssI*) genes (53). VgrG encodes a spike protein that, together with the PAAR adaptor protein, forms a membrane-penetrating needle (38,55), where different VgrG and PAAR variants typically associate with different types of effector molecules (31). We evaluated the presence of these three *vgrG* variants and found a large variability in their distribution across ST131 genomes (Fig. 5B). While different combinations occurred in variable proportions across niches, both *Poultry* and *Food* displayed a pattern that deviated from the rest, containing an enrichment of VgrG1 by itself (Fig. 5B).

**Figure 5.**
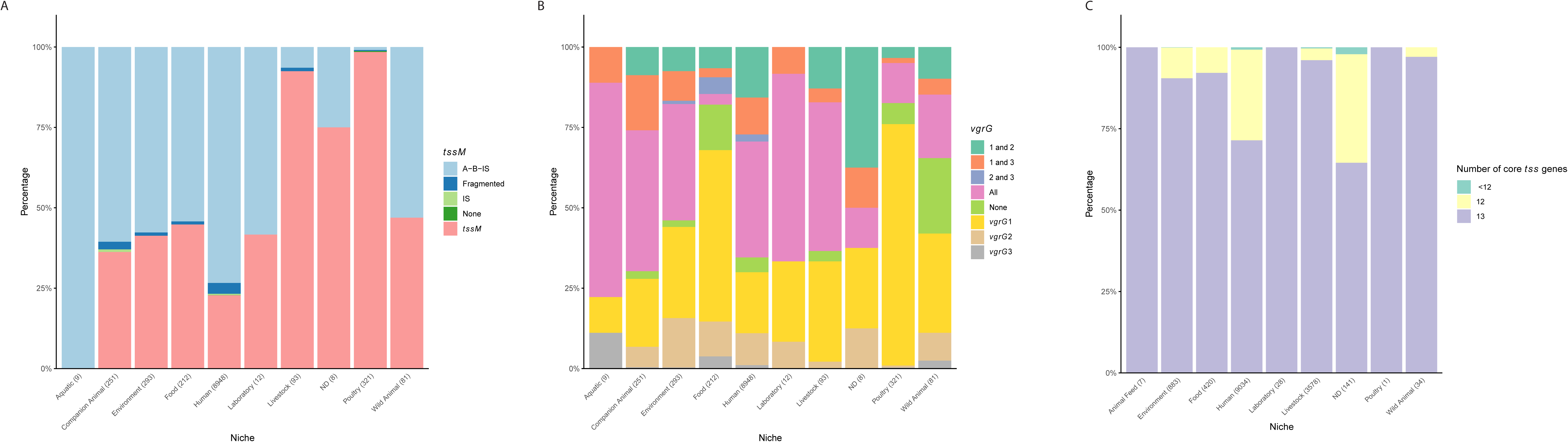
Niche-related T6SS patterns for ST131 and ST11. Calculated p-values to highlight which *Niche* groups differ significantly in their *Completeness* of T6SS^i2^ (for ST131) or T6SS^i1^ (for ST11) are listed in Table S8. **A.** Stacked bar graphs showing the relative distribution of T6SS^i2^-related *tssM* or its fragments in different niches. *A-B-IS* corresponds to the described fragmentation by IS*Ec12*(53) and detection of both fragments (A, B) and the IS, *Fragmented* corresponds to detection of other combinations of fragments (only fragment A or B, both fragments without IS, fragment A and IS, or fragment B and IS), *IS* to detection of only the IS, *None* to no detection of any of the elements, and *tssM* to the detection of an intact *tssM* gene. To provide broader context of *tssM* fragmentation, the same figure for the whole collection of genomes is shown in Fig. S10, which also gives an overview across phylogenetic groups. **B.** Stacked bar graphs showing the relative distributions of ST131-relevant *vgrG* (here referred to as 1-3) in different niches for ST131. The same figure for the whole collection of genomes is provided in Fig. S10, which also provides an overview across phylogenetic groups. **C.** Stacked bar graphs showing the *Completeness* of T6SS^i1^ in ST11 in relation to *Niche*.

In parallel, we examined the T6SSs of *E. coli* ST11, the most dominant IPEC and the most represented ST in our collection (14126 genomes). For this ST, we observed a drop in T6SS^i1^ *Completeness* (Fig. 4), which was linked to the disappearance of *vgrG*. This disappearance also displayed a niche-related pattern, occurring more in strains from the *Human* category than in *Livestock* (p-value < 10^-200^; Fig. 5C), the other dominant category. However, given that *vgrG* is a versatile T6SS component that is often located in an orphan region, it is less likely that this drop indeed corresponds to a loss of T6SS functionality. The drop could thus simply reflect the presence of an alternative VgrG variant, not present in our T6SS database, which occurs more in genomes from a *Human* source.

### T6SS co-occurrence analysis with virulence-associated genes and multi-drug resistance

In a final step, we sought to investigate the co-occurrence of T6SSs with other VAGs and ARGs, given that links between these and T6SSs have been described before (53,56) and could be important in a pathogenic context. For this, we examined the correlation between the *Presence* and *Completeness* of T6SS subclasses, VAGs, and multi-drug resistance (MDR; in this case determined by specified amounts of ARGs). The results of this analysis are summarized in the clustergram of Figure 6 and Table S9, and revealed two dominant clusters within the *E. coli* VAGs, which largely corresponded to IPEC and ExPEC-associated virulence factors. In line with our previous results based on the clinical associations and phylogeny of genomes (Fig. 2 and 3), T6SS^i1^ clustered with IPEC VAGs, while T6SS^i2^ clustered with ExPEC VAGs (Fig. 6A). Within the ExPEC-associated VAGs, we could identify several smaller subclusters that displayed strong correlations, indicating that these sets of VAGs typically co-occur (Fig. 6A, Cluster 1). We did not observe such smaller clusters for IPEC-associated VAGs, which mostly all co-occurred, with exception of colonization factor antigen I represented by *cfaA* (Fig. 6A, Cluster 2). In general, we found that most VAGs displayed mild positive correlations with one T6SS^i^ subclass, and negative for the other, with the low prevalence of T6SS^i4^ precluding a reliable analysis for this subclass.

**Figure 6:**
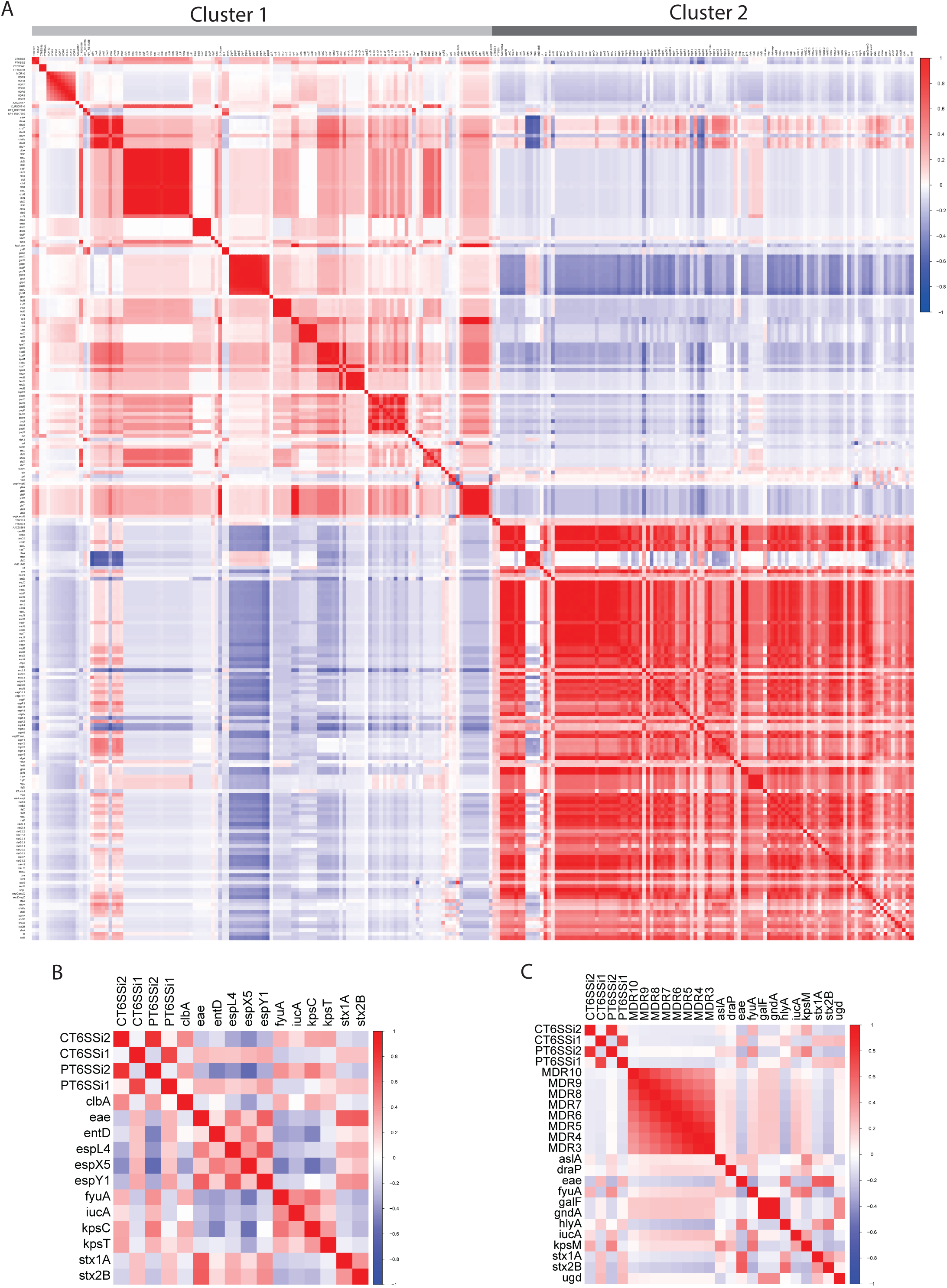
Correlations between T6SSi, VAGs and MDR. **A.** Clustergram depicting the correlation coefficients between the *Presence* and *Completeness* of T6SS^i^ subclasses, a collection of 227 VAGs recovered from VFDB, and multi-drug resistance. Cluster 1 contains typical ExPEC VAGs, while Cluster 2 contains typical IPEC VAGs. Fig. S11 provides a zoom in into the separate clusters. **B.** Heatmap highlighting the subset of individual VAGs displaying strong positive and negative correlations between *Presence* and *Completeness* of T6SS^i^ subclasses. **C.** Heatmap highlighting correlations between VAGs or T6SS^i^ subclasses and MDR.

While no correlations between the *Presence* and *Completeness* of different T6SSs and individual VAGs were extreme (Fig. 6 and Table S9), we did find some notable positive and negative ones (Fig. 6B). The highest positive correlation for T6SS^i1^ was with *espY1*, which is linked to Type III secretion system (T3SS) effectors (Lan and Shi, 2022) (57). T6SS^i1^ correlated positively with enterobactin (*entD*) (58) and shiga-like toxins (*stx2B*, *stx1A*, *eae)* (2), and negatively with siderophores including yersiniabactin and aerobactin *iucA*. On the other hand, T6SS^i2^ showed a negative correlation to other types of T3SS effectors – *espL4* and *espX5* (57), and was positively correlated with yersiniabactin represented by *fyuA* (59) and colibactin (*clbA*). T6SS^i2^ was also correlated positively with capsule-related genes such as *kpsC* and *kpsT* (60).

For MDR, we found a mild negative correlation with *Complete* T6SS^i1^ and T6SS^i2^ (Fig. 6C), for all levels of MDR definition (eight groups, ranging from minimally three to minimally ten ARGs; see also Methods). In general, MDR also appeared to co-occur more with ExPEC-associated VAGs than with IPEC ones. Mild positive correlations with MDR were observed mostly for genes related to the capsule such as *kpsM*, *galF*, *gndA* or *ugd* (60,61), yersiniabactin, aerobactin, adhesion (*draP*) or invasion (*aslA*). Mild negative correlations were observed for IPEC-associated shiga-like toxins. Interestingly, the hemolysin A (*hlyA*) correlated mildly but most negatively from all the VAGs with MDR. This exotoxin is considered one of the typical ExPEC-associated markers (2) and has been proposed as a high priority target for ExPEC vaccine development (62). However, our analysis showed that while it can be detected prevalently in some dominant ExPEC STs (ST73, ST127, ST372 and ST12), it is more often associated with dominant IPEC STs (ST11, ST17, ST21, ST32, ST33, ST16, ST29, ST442 and ST655) (Fig. S1C and Table 1) and overall, it is present in 22% of ExPEC and 71% of IPEC.

## Discussion

In this work, we compiled a large-scale *E. coli* genome collection to assess the distribution of T6SS subclasses across *E. coli*, and investigate their potential association with pathogenicity, virulence factors, and antibiotic resistance. Our analysis highlights both the extensive genomic diversity and the variable distribution of different T6SSs across *E. coli*.

By screening the *E. coli* genomes and their associated metadata, we subdivided the genomes into eight categories based on their clinical relevance/information (*Source curated*). This division revealed that the majority of *ExPEC* genomes belonged to phylogroups B2, D and F (Fig. 1B and 3), which is consistent with previous research (2,7,28). Also in line with previous studies (20) was the fact that we found most *IPEC* genomes in phylogroups B1 and E (Fig. 1B). While the latter PG has the highest relative association with *IPEC* and contained the most dominant IPEC ST (i.e., ST11, representing approx. one third of all IPEC strains; Fig. 4), the former contained almost the same absolute amount of *IPEC* genomes and includes notable IPEC STs such as ST21 and ST17 (Fig. 4). The fourth most dominant IPEC ST, ST10 (a dominant EPEC pathovar) is also commonly detected as a prominent ExPEC ST (15). In our collection, this ST was the third most common (7513 genomes, Table 2), with only a small proportion linked to IPEC (1239 genomes) or ExPEC (480 genomes) (Figure S9). An explanation for this observation could stem from this ST containing 0 ST-specific core genes in its pangenome (4), which indicates a lack of shared accessory genome within this ST in addition to the general *E. coli* core genome. Therefore, it is likely that this generalist lineage, which is often commensal (4), can possess specific accessory genes that enable differentiation into IPEC, ExPEC or hybrid IPEC/ExPEC pathogroups. While the majority of strains fall into a single category (either IPEC or ExPEC), reports of a hybrid aEPEC/ExPEC pathotype of ST10 do exist (63).

While we largely relied on the established virulence scheme to assign an *IPEC* label to strains, this was different for the *ExPEC* label. Given that virulence-based schemes have proven unreliable to identify ExPEC (2,15), we opted for a metadata-based approach. For the further subdivision into ExPEC pathovars, we used the classical ExPEC labels such as *UPEC*, *BSI* (classically named SEPEC), and *APEC*, but also added a *Respiratory* group. We added this last category as *E. coli* has the potential to cause respiratory infections, albeit with low incidence (64), which is also confirmed by our own analysis (407 out of 20523 ExPEC genomes). For UPEC- and BSI-related genomes, the most dominant STs were identical (i.e., ST131, followed by ST73, ST69, ST95 and ST1193) (Table S4), which aligns with previous reports of the potential of urinary-tract infections to progress into blood-stream infections and cause sepsis (65,66). Within the 35 most prevalent STs of *E. coli* that we identified (Fig. 4), we generally found a good agreement between the our ExPEC assignment and that of a previous study (15). The most notable difference was ST372, which was previously not identified as a dominant ExPEC ST, yet contained 480 *ExPEC* genomes in our analysis. This discrepancy could be explained by the strong association of this ST with *Companion animals*, causing it to be overlooked. At the same time, it is important to acknowledge that our metadata-based assignment of categories also has limitations. For example, we found that PG G was not strongly associated with ExPEC, in contrast to previous reports (21). However, this is likely a consequence of its link with avian, and especially poultry, sources, which are less likely to have ExPEC-relevant metadata reported. In addition, the assignment is also not straightforward due to potential secondary contamination of poultry body parts in slaughter houses. Similarly, we did not find a large representation of the APEC pathotype (446 genomes) across our collection, which likely reflects the same constraints of our approach (rather than APEC truly being a rare pathotype).

For the examination of the distribution of T6SS subclasses across *E. coli*, we made a distinction between *Present* (i.e., detection of at least 12 core genes) and *Complete* (i.e., detection of all 13 core genes). While *Present* can be seen as a liberal approximation of functionality, it likely represents an overestimation as the interruption of a single core gene has been reported before (53) and is something we also observed in our analysis (Fig. 4). *Complete*, on the other hand, likely represents an underestimation of functionality. This is an inherent consequence of working with databases of T6SS with confirmed functionality, which will never be complete. As a result, we expect the true levels of functional T6SS to lie somewhere between the reported prevalences of *Complete* and *Present*. Importantly, as we used a bioinformatics-based approach, the functionality can only be presumed, not confirmed, especially considering the thresholds we used for query coverage and identity (85 %). Despite these limitations, our approach enabled us to map the prevalence, distribution and diversity of T6SSs across *E. coli* at an unprecedented scale.

Our general analysis revealed a peculiar distribution pattern of T6SS^i^ subclasses across *E. coli*, where T6SS^i1^ was broadly distributed across all PGs, T6SS^i2^ was mostly associated with PGs B1, B2, and G, and T6SS^i4b^ was only found a small fraction of genomes belonging to PGs A, B1, and D (Fig. 2C). The distribution pattern of T6SS^i2^ did not correspond to that of a previous study, which indicated a main association with PG D. However, two main factors could underlie this discrepancy. First, Ma et al. (47) focused on T6SSs solely in APEC and performed their screen on a much smaller scale (472 APEC genomes, of which only 69 contained a T6SS^i2^). Despite this limitation, their findings were taken over and extrapolated across *E. coli* in reviews and research articles concerning T6SS in *E. coli* (38,46,48). As such, our findings reveal the importance of performing large-scale studies before extrapolating certain findings to a larger group of strains. Second, PG assignment has evolved over time. For example, ST117 is one of the most prevalent APEC STs (21,67) (also the most prevalent in this study) and is currently assigned to the relatively recent PG G, in which it represents the vast majority of strains (54). However, in 2013, the assignment, using the previous scheme (68), was PG D. Therefore, it is possible that the evolving PG scheme underlies at least part of the observed discrepancies. As such, we believe it is crucial to always confirm the correct PG classification, especially given that incorrect assignments tend to persist over time and lead to confusion (e.g., ST117 can be still found reported as PG D (69) or as PG F (53) in the recent literature).

Closer examination of T6SSs in ST131, the most notable ExPEC ST, revealed a number of potentially relevant properties in terms of its reduced *Completeness* (Fig. 5A). While this was due to a *tssM* fragmentation in ST131, which had been reported before (53), we also observed a clear niche-related pattern of this fragmentation (Fig. 5A). The disruption was most prevalent in *Humans*, yet almost absent in *Poultry* and *Livestock*. While this observation challenges the notion that ExPEC clones can be easily exchanged among these niches, it aligns with another study that showed that avian ST131 formed a distant cluster with specific VAGs (70). At the same time, this truncation could provide valuable information on the dominance of ST131 within ExPEC infections, given that most other ExPEC STs from the B2 phylogroup still contained a *Complete* T6SS^i2^ (Fig. 4). This observation suggests that T6SS^i2^ potentially contributes to the initial evolution of successful ExPEC lineages, and that its subsequent loss of functionality confers additional cellular properties relevant for the pathogenicity of ST131. These properties currently remain speculative but could include enhanced fitness and stress resistance (as reported for *Campylobacter jejuni* in the presence of bile salts (71)), or enhanced horizontal gene transfer. Horizontal gene transfer is known to allow *E. coli* adapt to new niches and the large accessory genome of ST131, consisting of over 22500 genes (72), suggest this is at least one of the factors ST131 has benefitted from. In addition, the propagation of conjugation has been linked to the repression of T6SSs (73), and ST131 without functional T6SS has also been linked to a higher level of multi-drug resistance which is likely connected with higher level of conjugation events (53). Our own analysis also indicated a mild negative association between functional T6SS^i2^ and MDR across *E. coli* genomes (Fig. 6). These two factors combined (i.e., increased stress resistance and MDR acquisition through enhanced horizontal gene transfer) could contribute the global pathogenic success of ST131.

Our co-occurrence analysis revealed that T6SS^i1^ mostly clustered with IPEC VAGs, while T6SS^i2^ largely clustered with ExPEC VAGs (Figure 6). T6SS^i1^ occurrence correlated positively with many typical IPEC VAGs such as *stx1* and *stx2*, or T3SS-related genes such as *escC* (2,5), and T6SS^i2^ occurrence correlated positively with typical ExPEC markers such as capsule or yersiniabactin (3). In general, we did not observe clustering of IPEC together with ExPEC VAGs. Their co-occurrence analysis also displayed negative correlations, indicating that these pathogroups are, in the majority of cases, mutually exclusive. This type of global patterns could only be identified through our merged database of *E. coli* T6SS^i^ regions that combines input from multiple sources, given that different sources showed different distribution patterns for T6SS subclasses.

In conclusion, this study provides a comprehensive overview on T6SS^i^ subclass distribution within *E. coli*, and examines the relationship of its presence with other clinically relevant information such as the source of isolation, the clinical context, VAGs, and MDR. Moreover, we are confident that our large-scale annotated *E. coli* genome collection will serve as a useful resource to examine the distribution and prevalence with regard to the clinical context of other genetic elements in the future.

## Materials and Methods

### Genomic data collection and annotation

We downloaded a total of 136051 available genomes with metadata from EB from February to April 2023. In addition, we downloaded three types of metadata datasets from EB, *Assembly stats*, *Phylotype*, and *Annotation*, and merged them using *Uberstrain* as a barcode to match these datasets. We then used an *Assembly barcode* (and relabeled it as *Barcode*) to match the metadata with the results of the evaluation of the genomic sequences. To determine phylogenetic groups, we used the ClermonTyping tool (https://github.com/A-BN/ClermonTyping), and we merged our results with EB metadata (EB also provides a phylogenetic group assignment based on either ClermonTyping (54) or EZClermont (74) in the *Phylotype* metadata sheet (27)). We filtered out genomes that showed irregular PG or species results (269 genomes), or were detected as part of very minor clades III, IV or V (total of 65 genomes). We filtered out additional genomes (4107) due to low coverage (below 30x), resulting in a final dataset of 131610 genomes.

Using the existing EB metadata, we provided additional annotations (Table S1) by creating four columns *Source curated*, *ExPEC type*, *IPEC type* and *Gender*, and updating information in the columns *Source Pathogen* (in EB originally called *Path/Nonpath*) and *Source Pathogen details* (originally *Simple Patho*) as these occasionally contain information not relevant for our purposes (e.g., AMR profile).

*Source curated* divided the genomes into eight categories: *Clinical* (1136 genomes), *ExPEC* (20523 genomes), *Healthy-Community* (571 genomes), *IPEC* (42791 genomes), *Mixed* (159 genomes), *ND* (Not determined) (65828 genomes), *Non Pathogen* (183 genomes), and *Pathogen* (419 genomes). The category *Clinical* contains genomes with keywords *Hospital*, *Clinical*, or *Patient* in their metadata, which thus likely correspond to pathogenic strains, but where we lack the necessary information to reliably divide them into the *IPEC* or *ExPEC* category. The *ExPEC* category was assigned based on existing metadata, if ExPEC was already specified in the *Source Pathogen* or *Source Pathogen* details, *Simple Disease* or *Disease* information, or if the source of isolation was ExPEC-relevant (e.g., urine, blood). For other sources (e.g., skin, eyes, ears), infection needed to be mentioned to assign the *ExPEC* label. The *Healthy-Community* category contains genomes for which it was specifically mentioned that they originated from healthy individuals (humans and animals) or from the community (“healthy” or “community”). The assignment of genomes to the *IPEC* category was mainly based on Pathovar predictions using the established virulence scheme (recovered from EB dataset *Phylotype*). A smaller part of IPEC assignment was based on the direct contributor’s metadata. The *Mixed* category contains genomes for which both the *ExPEC* and *IPEC* label would be applicable (e.g., a genome with predicted IPEC pathovar isolated from an urinary infection). Genomes for which further annotation was not possible based on the provided metadata were classified as *ND*. The *Non Pathogen* category contains genomes for which this information was specified by the EB contributors (“non-pathogen”) and no other apparently contradicting metadata exists. Similarly, the *Pathogen* category contains genomes for which this information was provided by the contributors (“pathogen”) and no other metadata could link these genomes to the IPEC or ExPEC pathogroup. For the *Clinical*, *ND*, and *Pathogen* category, we expect pathogenic strains to be enriched in ExPEC genomes, given that IPEC can be identified based on their genomic information and a well-defined VAGs scheme.

The columns *ExPEC type* and *IPEC type* (Table S1) further divide these two pathogroups into specific pathovars or other defined groups. In the case of ExPEC, we deviated from the classically defined pathovars due to unreliability of the assignment and an underrepresentation in some cases (e.g., neonatal meningitis-causing *E. coli* (NMEC) was specified for only four strains in the collection and only 26 additional strains were linked to the brain). Our division of *ExPEC type* contained five categories: *UPEC* (12346 genomes, linked to urine or UTIs), *BSI* (4957 genomes, linked to blood), *Respiratory* (407 genomes, linked to respiratory infection or intubation), *APEC* (446 genomes, linked to avian ExPEC) and other unspecified *ExPEC* (2524 genomes, other sources). If there was not sufficient metadata to reliably determine whether a genome was ExPEC-linked or not, genomes were assigned to the category *NS* (68322 genomes). If a genome was already linked to an IPEC pathotype, and thus likely not ExPEC, it was assigned to the category *No* (42608 genomes). *IPEC type* was, in the majority of cases (> 95%), assigned based on the virulence scheme of genomes (Table S1). The pathovars included *EHEC* (24298 genomes), *STEC* (9966 genomes), *EPEC* (5176 genomes), *ETEC* (2266 genomes), *EIEC* (356 genomes) and hybrid groups of *ETEC/STEC* (557 genomes), *ETEC/EHEC* (102 genomes) and *ETEC/EPEC* (46 genomes). All remaining strains for which the virulence scheme showed negative results were assigned into the *No* category (88342 genomes), with exception of strains for which the virulence scheme results were missing, which are grouped in the *ND* category (501 genomes). The column *Gender* is specified in the rare instances where the genome originated from a female- or male-specific source (e.g., vagina, uterus, semen) or when it was specifically mentioned by the EB contributors.

Further metadata adjustments included adding missing *Continent* info base on *Country* and adding *Source Niche* data where possible from *Source type* or *Source details*. There were only two genomes with original *Continent* label *Central America* (Uberstrains ESC_GA0177AA and ESC_GA0176AA), which were merged with *North America*. For *Source Niche*, *Aquatic Animal* (603 genomes) was merged with *Aquatic* (1 genome, Uberstrain ESC_RA0094AA, which also came from an aquatic animal). The merged group was named *Aquatic*.

### Screen for T6SS components

To construct a comprehensive sequence database of *E. coli* T6SS components, we used multiple sources as genomic references of T6SS components of different T6SS classes and subclasses. We always used ABRicate (https://github.com/tseemann/abricate) to screen for the presence of T6SS-associated genes with a threshold of 85% for both query coverage and identity. A first source came from a screen for ExPEC-associated regions, where we identified a T6SS in a strain of ST131 (KO_178_B, originally isolated in Tausova *et al*., 2012, BioSample SAMN35728132) (75). This same region was identified in a previous study (53), and showed only one SNP difference with ours. To investigate the broader occurrence of various T6SS^i^ subclasses, we downloaded sequences of ten T6SS regions identified by Ma et al. (47), who identified the *E. coli*-related subclasses T6SS^i1^, T6SS^i2^ and T6SS^i4b^ across APEC genomes. When necessary, we provided additional annotations to assign genes as *tssA*-*tssM* using BLAST (76) or the SecReT6 database (https://bioinfo-mml.sjtu.edu.cn/SecReT6/index.php) (77) (Table S2). Given that this approach did not always find all 13 core components within identified T6SS genomic regions, we also implemented all reported functional *E. coli* T6SS regions from the SecReT6 database. We evaluated both the completeness and presence of the three T6SS subclasses. We defined *Present/Presence* (P) as detecting at least 12 core genes from a respective subclass, while *Complete/Completeness* (C) corresponds to the detection of all 13 core genes. We performed this analysis using our merged database, but also separately using only the sequences from Ma & KO_178_B (DB1) or SecReT6 (DB2) (Fig. S3-7).

### Phylogeny of the most prevalent ST representatives

Across our genome collection, we identified 35 prevalent STs (using a cut-off of ≥ 500 genomes for a given ST). We pseudo-randomly selected one genome per ST (i.e., typically one of the first genomes per ST in alphabetical order by *Assembly barcode* with the intention to include diverse countries and sources of origin) and used it to build a phylogenetic tree to visualize the prevalence of T6SSs in the approximate phylogenetic context. The genomes used to build the tree are specified in Fig. S1. The phylogenetic analysis was based on Prokka (v1.14.5) open reading frames prediction (78), and Roary (v3.7.0) (79) multi-fasta alignment RAxML (80), supported by 100 bootstraps, was used to calculate the tree. The prevalence of individual T6SS^i^ subclasses was examined in terms of both their *Presenc*e and *Completeness*. In addition, we separately inspected STs with large differences between both categories by checking the prevalence of individual genes and identifying those that were missing (and thus responsible for the discrepancy between *Presence* and *Completeness*). For each such ST, we examined an individual genome sequence and visually inspected the T6SS region to uncover the potential reason underlying the disappearance of the individual gene.

### Virulence Factor Databese (VFDB) and ResFinder screen

We used the VFDB (http://www.mgc.ac.cn/VFs/) (61) to screen for the presence of known VAGs in our *E. coli* genome collection using ABRicate with a 90% threshold for both query coverage and identity. We compared the obtained list with the distribution pattern of T6SS and examined potential correlations in their mutual presence or absence. To obtain meaningful results, we limited this screen to VAGs that are found in more than 3% and less than 90% of genomes in our collection. This resulted in a list of 242 VAGs (from a total of 404 that were originally identified). The prevalence of these 242 VAGs for the 35 dominant STs is summarized in Fig. S1. For our correlation analysis, we filtered out T6SS-relevant genes within the VFDB, which were found to mostly associate with T6SS^i1^, to avoid artificial positive correlations. This resulted in a total of 227 VAGs that were considered for the global correlation analysis. To examine the potential co-occurrence of T6SS with MDR, we used ResFinder (v.3.0) (81) to screen for the presence of ARGs. We filtered out *mdf(A)*, given that it is normally present in *E. coli* (82,83) and then we calculated the sum of overall detected ARGs. We defined the MDR on multiple levels, starting from genomes with minimally three ARGs, and ending with genomes with minimally ten ARGs, yielding 8 MDR groups in total, marked MDR3-MDR10. The correlations of presence/absence of T6SS, VAGs and MDR were calculated using polychoric correlation coefficients, the code for this is reachable at https://github.com/Govers-Lab/Nesporova_and_Govers_2024.

### Statistical analysis and data visualization

We used RStudio (v2023.06.0) with R (v4.4.1) for data merge, summary, statistical analysis and figure creation. The code is specified and explained at https://github.com/Govers-Lab/Nesporova_and_Govers_2024 together with raw and intermediate data. We used pairwise Wilcoxon test to calculate the significance (α = 0.05) of the pairwise differences between specific groups with Benjamini-Hochberg adjustment of the p-values. We used Geneious (v7.1.9) to check annotations and alignment or reference sequences and specific genomic regions. The phylogenetic tree was visualized using the Interactive Tree of Life (iTOL v6) (84).

### Availability of data and materials

All data generated or analyzed during this study are included in this published article and it supplementary information files.

## Acknowledgments

This work was funded by the European Union under Horizon Europe: MSCA Actions (Project 101105027) and by junior postdoctoral fellowship from the Research Foundation – Flanders (file number 1251624N) awarded to K.N. This work was further supported by a start-up grant (STG/21/068) from the KU Leuven Research Fund awarded to S.K.G. We would also like to thank Prof. Monika Dolejska for providing the long-read sequence for KO_178_B and the members of Govers laboratory for valuable feedback on the manuscript. We acknowledge all EnteroBase contributors.

## Author contributions

Conceptualization, K.N. and S.K.G.; methodology, K.N. and S.K.G.; software, K.N.; formal analysis, K.N.; investigation, K.N.; data curation, K.N.; writing – original draft, K.N. and S.K.G.; writing – review & editing, K.N. and S.K.G.; visualization, K.N. and S.K.G.; supervision, S.K.G.; project administration, S.K.G.; funding acquisition, K.N. and S.K.G.

## References

1. Martinson JNV, Walk ST. 2020. *Escherichia coli* Residency in the Gut of Healthy Human Adults. EcoSal Plus 9(1).

2. Geurtsen J, de Been M, Weerdenburg E, Zomer A, McNally A, Poolman J. 2022. Genomics and pathotypes of the many faces of *Escherichia coli*. FEMS Microbiol Rev 46(6).

3. Dale AP, Woodford N. 2015. Extra-intestinal pathogenic *Escherichia coli* (ExPEC): Disease, carriage and clones. J Infect 71(6):615–26.

4. Cummins EA, Hall RJ, Connor C, McInerney JO, McNally A. 2022. Distinct evolutionary trajectories in the *Escherichia coli* pangenome occur within sequence types. Microb Genomics 8(11)mgen000903.

5. Croxen MA, Law RJ, Scholz R, Keeney KM, Wlodarska M, Finlay BB. 2013. Recent advances in understanding enteric pathogenic *Escherichia coli*. Clin Microbiol Rev 26(4):822–80.

6. Poirel L, Madec J-Y, Lupo A, Schink A-K, Kieffer N, Nordmann P, Schwarz S. 2018. Antimicrobial Resistance in *Escherichia coli*. Microbiol Spectr 6(4):10.1128/microbiolspec.arba-0026-2017.

7. McNally A, Kallonen T, Connor C, Abudahab K, Aanensen DM, Horner C, Peacock SJ, Parkhill J, Croucher NJ, Corander J. 2019. Diversification of colonization factors in a multidrug-resistant *Escherichia coli* lineage evolving under negative frequency-dependent selection. mBio 10(2):e00644–19.

8. Laupland KB, Church DL. 2024. Population-based epidemiology and microbiology of community-onset bloodstream infections. Clin Microbiol Rev 27(4):647–64.

9. Riley LW. 2020. Distinguishing Pathovars from Nonpathovars: *Escherichia coli*. Microbiol Spectr 8(4):10.1128/microbiolspec.ame-0014-2020.

10. Kaper JB, Nataro JP, Mobley HLT. 2004. Pathogenic *Escherichia coli*. Nat Rev Microbiol 2(2):123–40.

11. Rasko DA, Webster DR, Sahl JW, Bashir A, Boisen N, Scheutz F, Paxinos EE, Sebra R, Chin CS, Iliopoulos D, Klammer A, Peluso P, Lee L, Kislyuk AO, Bullard J, Kasarskis A, Wang S, Eid J, Rank D, Redman JC, Steyert SR, Frimodt-Møller J, Struve C, Petersen AM, Krogfelt KA, Nataro JP, Schadt EE, Waldor MK. 2011. Origins of the *E. coli* strain causing an outbreak of hemolytic-uremic syndrome in Germany. N Engl J Med 365(8):709–17.

12. Li D, Reid CJ, Kudinha T, Jarocki VM, Djordjevic SP. 2020. Genomic analysis of trimethoprim-resistant extraintestinal pathogenic *Escherichia coli* and recurrent urinary tract infections. Microb Genomics 6(12):mgen000475.

13. Clermont O, Christenson JK, Denamur E, Gordon DM. 2013. The Clermont *Escherichia coli* phylo-typing method revisited: improvement of specificity and detection of new phylo-groups. Environ Microbiol Rep 5(1):58–65.

14. Wirth T, Falush D, Lan R, Colles F, Mensa P, Wieler LH, Karch H, Reeves PR, Maiden MC, Ochman H, Achtman M. 2006. Sex and virulence in *Escherichia coli*: An evolutionary perspective. Mol Microbiol 60(5):1136–51.

15. Manges AR, Geum HM, Guo A, Edens TJ, Fibke CD, Pitout JDD. 2019. Global Extraintestinal Pathogenic *Escherichia coli* (ExPEC) Lineages. Clin Microbiol Rev 12;32(3):e00135–18.

16. Scallan E, Hoekstra RM, Angulo FJ, Tauxe R V., Widdowson MA, Roy SL, Jones JL, Griffin PM. 2011. Foodborne illness acquired in the United States-Major pathogens. Emerg Infect Dis 17(1):7–15.

17. Rangel JM, Sparling PH, Crowe C, Griffin PM, Swerdlow DL. 2005. Epidemiology of *Escherichia coli* O157:H7 outbreaks, United States, 1982-2002. Emerg Infect Dis 11(4):603–9.

18. Russo TA, Johnson JR. 2000. Proposal for a new inclusive designation for extraintestinal pathogenic isolates of *Escherichia coli*: ExPEC. J Infect Dis 181(5):1753–4.

19. Russo TA, Johnson JR. 2003. Medical and economic impact of extraintestinal infections due to *Escherichia coli*: Focus on an increasingly important endemic problem. Microbes Infect 5(5):449–56.

20. Denamur E, Clermont O, Bonacorsi S, Gordon D. 2021. The population genetics of pathogenic *Escherichia coli*. Nat Rev Microbiol 19(1):37–54.

21. Mageiros L, Méric G, Bayliss SC, Pensar J, Pascoe B, Mourkas E, Calland JK, Yahara K, Murray S, Wilkinson TS, Williams LK, Hitchings MD, Porter J, Kemmett K, Feil EJ, Jolley KA, Williams NJ, Corander J, Sheppard SK. 2021. Genome evolution and the emergence of pathogenicity in avian *Escherichia coli*. Nat Commun 22;12(1):1934.

22. Djordjevic SP, Jarocki VM, Seemann T, Cummins ML, Watt AE, Drigo B, Wyrsch ER, Reid CJ, Donner E, Howden BP. 2024. Genomic surveillance for antimicrobial resistance — a One Health perspective. Nat Rev Genet 25(2):142–57.

23. Stein RA, Katz DE. 2017. *Escherichia coli*, cattle and the propagation of disease. FEMS Microbiol Lett 364(6):fnx050.

24. Weiner-Lastinger LM, Abner S, Benin AL, Edwards JR, Kallen AJ, Karlsson M, Magill SS, Pollock D, See I, Soe MM, Walters MS, Dudeck MA. 2020. Antimicrobial-resistant pathogens associated with pediatric healthcare-associated infections: Summary of data reported to the National Healthcare Safety Network, 2015-2017. Infect Control Hosp Epidemiol 41(1):19–30.

25. Fay K, Sapiano MRP, Gokhale R, Dantes R, Thompson N, Katz DE, Ray SM, Wilson LE, Perlmutter R, Nadle J, Godine D, Frank L, Brousseau G, Johnston H, Bamberg W, Dumyati G, Nelson D, Lynfield R, DeSilva M, Kainer M, Zhang A, Ocampo V, Samper M, Pierce R, Irizarry L, Sievers M, Maloney M, Fiore A, Magill SS, Epstein L. 2020. Assessment of Health Care Exposures and Outcomes in Adult Patients with Sepsis and Septic Shock. JAMA Netw Open 3(7):e206004.

26. Rhee C, Klompas M. 2020. Sepsis trends: Increasing incidence and decreasing mortality, or changing denominator? J Thorac Dis 2(Suppl 1):S89–100.

27. Zhou Z, Alikhan NF, Mohamed K, Fan Y, Achtman M. 2020. The EnteroBase user’s guide, with case studies on Salmonella transmissions, *Yersinia pestis* phylogeny, and *Escherichia* core genomic diversity. Genome Res 30(1):138–52.

28. Abram K, Udaondo Z, Bleker C, Wanchai V, Wassenaar TM, Robeson MS, Robeson MS 2nd, Ussery DW. 2021. Mash-based analyses of *Escherichia coli* genomes reveal 14 distinct phylogroups. Commun Biol 4(1):117.

29. Reid CJ, Cummins ML, Börjesson S, Brouwer MSM, Hasman H, Hammerum AM, Roer L, Hess S, Berendonk T, Nesporova K, Haenni M, Madec JY, Bethe A, Michael GB, Schink AK, Schwarz S, Dolejska M, Djordjevic SP. 2022. A role for ColV plasmids in the evolution of pathogenic *Escherichia coli* ST58. Nat Commun 13(1):683.

30. Allsopp LP, Bernal P, Nolan LM, Filloux A. 2020. Causalities of war: The connection between type VI secretion system and microbiota. Cell Microbiol 22(3):e13153.

31. Geller AM, Shalom M, Zlotkin D, Blum N, Levy A. 2024. Identification of type VI secretion system effector-immunity pairs using structural bioinformatics. Mol Syst Biol 20(6):702–18.

32. Unni R, Pintor KL, Diepold A, Unterweger D. 2022. Presence and absence of type VI secretion systems in bacteria. Microbiol (United Kingdom) 168(4).

33. Singh RP, Kumari K. 2023. Bacterial type VI secretion system (T6SS): an evolved molecular weapon with diverse functionality. Biotechnol Lett 45(3):309–31.

34. Speare L, Cecere AG, Guckes KR, Smith S, Wollenberg MS, Mandel MJ, Miyashiro T, Septer AN. 2018. Bacterial symbionts use a type VI secretion system to eliminate competitors in their natural host. Proc Natl Acad Sci U S A 115(36):E8528–37.

35. Logan SL, Thomas J, Yan J, Baker RP, Shields DS, Xavier JB, Hammer BK, Parthasarathy R. 2018. The *Vibrio cholerae* type VI secretion system can modulate host intestinal mechanics to displace gut bacterial symbionts. Proc Natl Acad Sci U S A 115(16):E3779–87.

36. Drebes Dörr NC, Blokesch M. 2018. Bacterial type VI secretion system facilitates niche domination. Proc Natl Acad Sci U S A 115(36):8855–7.

37. Pukatzki S, Ma AT, Revel AT, Sturtevant D, Mekalanos JJ.2007. Type VI secretion system translocates a phage tail spike-like protein into target cells where it cross-links actin. Proc Natl Acad Sci U S A 104(39):15508–13.

38. Journet L, Cascales E. 2016. The Type VI Secretion System in *Escherichia coli* and Related Species . EcoSal Plus 7(1):10.1128/ecosalplus.ESP-0009-2015.

39. Mariano G, Trunk K, Williams DJ, Monlezun L, Strahl H, Pitt SJ, Coulthurst SJ. 2019. A family of Type VI secretion system effector proteins that form ion-selective pores. Nat Commun 10(1).

40. Zong B, Zhang Y, Wang X, Liu M, Zhang T, Zhu Y, Zheng Y, Hu L, Li P, Chen H, Tan C. 2019. Characterization of multiple type-VI secretion system (T6SS) VgrG proteins in the pathogenicity and antibacterial activity of porcine extra-intestinal pathogenic *Escherichia coli*. Virulence 10(1):118–32.

41. Robinson L, Liaw J, Omole Z, Xia D, van Vliet AHM, Corcionivoschi N, Hachani A, Gundogdu O. 2021. Bioinformatic Analysis of the *Campylobacter jejuni* Type VI Secretion System and Effector Prediction. Front Microbiol 12:694824.

42. Robinson LA, Collins ACZ, Murphy RA, Davies JC, Allsopp LP. 2023. Diversity and prevalence of type VI secretion system effectors in clinical *Pseudomonas aeruginosa* isolates. Front Microbiol 13:1042505.

43. Sana TG, Voulhoux R, Monack DM, Ize B, Bleves S. 2020. Protein Export and Secretion Among Bacterial Pathogens. Front Cell Infect Microbiol 9:473.

44. Gallegos-Monterrosa R, Coulthurst SJ. 2021. The ecological impact of a bacterial weapon: Microbial interactions and the Type VI secretion system. FEMS Microbiol Rev 45(6):fuab033.

45. Hu J, Afayibo DJA, Zhang B, Zhu H, Yao L, Guo W, Wang X, Wang Z, Wang D, Peng H, Tian M, Qi J, Wang S. 2022. Characteristics, pathogenic mechanism, zoonotic potential, drug resistance, and prevention of avian pathogenic *Escherichia coli* (APEC). Front Microbiol 13:1049391.

46. Navarro-Garcia F, Ruiz-Perez F, Cataldi Á, Larzábal M. 2019. Type VI secretion system in pathogenic *Escherichia coli*: Structure, role in virulence, and acquisition. Front Microbiol 10:1965.

47. Ma J, Sun M, Bao Y, Pan ZH, Zhang W, Lu C, Yao H. 2013. Genetic diversity and features analysis of type VI secretion systems loci in avian pathogenic *Escherichia coli* by wide genomic scanning. Infect Genet Evol 20:454–64.

48. Chen X, Liu W, Li H, Yan S, Jiang F, Cai W, Li G. 2021. Whole genome sequencing analysis of avian pathogenic *Escherichia coli* from China. Vet Microbiol 259:109158.

49. Micenková L, Frankovičová L, Jaborníková I, Bosák J, Dítě P, Šmarda J, Vrba M, Ševčíková A, Kmeťová M, Šmajs D. 2018. *Escherichia coli* isolates from patients with inflammatory bowel disease: ExPEC virulence- and colicin-determinants are more frequent compared to healthy controls. Int J Med Microbiol 308(5):498–504.

50. He W, Wu K, Ouyang Z, Bai Y, Luo W, Wu D, An H, Guo Y, Jiao M, Qin Q, Zhang J, Wu Y, She J, Hwang PM, Zheng F, Zhu L, Wen Y. 2023 Structure and assembly of type VI secretion system cargo delivery vehicle. Cell Rep 42(7):112781.

51. Clemens DL, Lee BY, Horwitz MA. 2018. The *Francisella* Type VI secretion system. Front Cell Infect Microbiol 8:121.

52. Lu L, Qi Z, Chen Z, Wang H, Wei X, Zhao B, Wang Z, Shao Y, Tu J, Song X. 2024. Avian pathogenic *Escherichia coli* T6SS effector protein Hcp2a causes mitochondrial dysfunction through interaction with LETM1 protein in DF-1 cells. Poult Sci 103(4):103514.

53. Cummins EA, Moran RA, Snaith AE, Hall RJ, Connor CH, Dunn SJ, McNally A. 2023. Parallel loss of type VI secretion systems in two multi-drug-resistant *Escherichia coli* lineages. Microb Genomics 9(11): 001133.

54. Beghain J, Bridier-Nahmias A, Nagard H Le, Denamur E, Clermont O. 2018. ClermonTyping: An easy-to-use and accurate in silico method for *Escherichia* genus strain phylotyping. Microb Genomics 4(7):e000192.

55. Douzi B, Brunet YR, Spinelli S, Lensi V, Legrand P., Blangy S, Kumar A, Journet L, Cascales E, Cambillau C. 2016. Structure and specificity of the Type VI secretion system ClpV-TssC interaction in enteroaggregative *Escherichia coli*. Sci Rep 6:34405.

56. Otto SB, Servajean R, Lemopoulos A, Bitbol AF, Blokesch M. 2024. Interactions between pili affect the outcome of bacterial competition driven by the type VI secretion system. Curr Biol 34(11):2403–2417.e9.

57. Lan L, Shi H. 2022. New insights into the formation and recovery of sublethally injured *Escherichia coli* O157:H7 induced by lactic acid. Food Microbiol 102:103918.

58. Martin P, Tronnet S, Garcie C, Oswald E. 2017. Interplay between siderophores and colibactin genotoxin in *Escherichia coli*. IUBMB Life 69(6):435–41.

59. Schubert S, Picard B, Gouriou S, Heesemann J, Denamur E. 2002. *Yersinia* high-pathogenicity island contributes to virulence in *Escherichia coli* causing extraintestinal infections. Infect Immun 70(9):5335–7.

60. Sande C, Whitfield C. 2021. Capsules and Extracellular Polysaccharides in *Escherichia coli* and Salmonella. EcoSal Plus 9(2):eESP00332020.

61. Chen L, Zheng D, Liu B, Yang J, Jin Q. 2016. VFDB 2016: Hierarchical and refined dataset for big data analysis - 10 years on. Nucleic Acids Res 44(D1):D694–7.

62. Xing Y, Clark JR, Chang JD, Zulk JJ, Chirman DM, Piedra F-A, Vaughan EE, Hernandez Santos HJ, Patras KA, Maresso AW. 2024. Progress toward a vaccine for extraintestinal pathogenic *E. coli* (ExPEC) II: efficacy of a toxin-autotransporter dual antigen approach. Infect Immun 7;92(5):e0044023.

63. Díaz-Jiménez D, García-Meniño I, Herrera A, Lestón L, Mora A. 2021. Microbiological risk assessment of Turkey and chicken meat for consumer: Significant differences regarding multidrug resistance, mcr or presence of hybrid aEPEC/ExPEC pathotypes of *E. coli*. Food Control 123.

64. Gong L, Tang N, Chen D, Sun K, Lan R, Zhang W, Zhou H, Yuan M, Chen X, Zhao X, Che J, Bai X, Zhang Y, Xu H, Walsh TR, Lu J, Xu J, Li J, Feng J. 2020. A Nosocomial Respiratory Infection Outbreak of Carbapenem-Resistant *Escherichia coli* ST131 With Multiple Transmissible *bla*_KPC–2_ Carrying Plasmids. Front Microbiol 11:2068.

65. Burgaya J, Marin J, Royer G, Condamine B, Gachet B, Clermont O, Jaureguy F, Burdet C, Lefort A, de Lastours V, Denamur E, Galardini M, Blanquart F; Colibafi/Septicoli & Coliville groups. 2023. The bacterial genetic determinants of *Escherichia coli* capacity to cause bloodstream infections in humans. PLoS Genet 19(8):e1010842.

66. Thänert R, Choi JH, Reske KA, Hink T, Thänert A, Wallace MA, Wang B, Seiler S, Cass C, Bost MH, Struttmann EL, Iqbal ZH, Sax SR, Fraser VJ, Baker AW, Foy KR, Williams B, Xu B, Capocci-Tolomeo P, Lautenbach E, Burnham CD, Dubberke ER, Kwon JH, Dantas G; CDC Prevention Epicenters Program. 2022. Persisting uropathogenic *Escherichia coli* lineages show signatures of niche-specific within-host adaptation mediated by mobile genetic elements. Cell Host Microbe 30(7):1034–1047.e6.

67. Mora A, López C, Herrera A, Viso S, Mamani R, Dhabi G, Alonso MP, Blanco M, Blanco JE, Blanco J. 2012. Emerging avian pathogenic *Escherichia coli* strains belonging to clonal groups O111: H4-D-ST2085 and O111: H4-D-ST117 with high virulence-gene content and zoonotic potential. Vet Microbiol 156(3–4):347–52.

68. Johnson JR, Owens KL, Clabots CR, Weissman SJ, Cannon SB. 2006. Phylogenetic relationships among clonal groups of extraintestinal pathogenic *Escherichia coli* as assessed by multi-locus sequence analysis. Microbes Infect 8(7):1702–13.

69. Belmahdi M, Chenouf NS, Ait Belkacem A, Martinez-Alvarez S, Pino-Hurtado MS, Benkhechiba Z, Lahrech S, Hakem A, Torres C. 2022. Extended Spectrum β-Lactamase-Producing *Escherichia coli* from Poultry and Wild Birds (Sparrow) in Djelfa (Algeria), with Frequent Detection of CTX-M-14 in Sparrow. Antibiotics 11(12):1814.

70. Bonnet R, Beyrouthy R, Haenni M, Nicolas-Chanoine MH, Dalmasso G, Madec JY. 2021. Host colonization as a major evolutionary force favoring the diversity and the emergence of the worldwide multidrug-resistant *Escherichia coli* ST131. MBio 12(4):e0145121.

71. Gupta S, Ray S, Khan A, China A, Das D, Mallick AI. 2021. The cost of bacterial predation via type VI secretion system leads to predator extinction under environmental stress. iScience 24(12):103507.

72. Decano AG, Downing T. 2019. An *Escherichia coli* ST131 pangenome atlas reveals population structure and evolution across 4,071 isolates. Sci Rep 9(1):17394.

73. Di Venanzio G, Moon KH, Weber BS, Lopez J, Ly PM, Potter RF, Dantas G, Feldman MF. 2019. Multidrug-resistant plasmids repress chromosomally encoded T6SS to enable their dissemination. Proc Natl Acad Sci U S A 22;116(4):1378–1383.

74. Waters NR, Abram F, Brennan F, Holmes A, Pritchard L. 2020. Easy phylotyping of *Escherichia coli* via the EzClermont web app and command-line tool. Access Microbiol 2(9):acmi000143.

75. Tausova D, Dolejska M, Cizek A, Hanusova L, Hrusakova J, Svoboda O, Camlik G, Literak I. 2012. *Escherichia coli* with extended-spectrum β-lactamase and plasmid-mediated quinolone resistance genes in great cormorants and mallards in Central Europe. J Antimicrob Chemother 67(5):1103–7.

76. Camacho C, Boratyn GM, Joukov V, Vera Alvarez R, Madden TL. 2023. ElasticBLAST: accelerating sequence search via cloud computing. BMC Bioinformatics 24(1):117.

77. Zhang J, Guan J, Wang M, Li G, Djordjevic M, Tai C, Wang H, Deng Z, Chen Z, Ou HY. 2023. SecReT6 update: a comprehensive resource of bacterial Type VI Secretion Systems. Sci China Life Sci 66(3):626–34.

78. Seemann T. 2014. Prokka: Rapid prokaryotic genome annotation. Bioinformatics. 30(14):2068–9.

79. Page AJ, Cummins CA, Hunt M, Wong VK, Reuter S, Holden MTG, Fookes M, Falush D, Keane JA, Parkhill J. 2015. Roary: Rapid large-scale prokaryote pan genome analysis. Bioinformatics 31(22):3691–3.

80. Stamatakis A. 2014. RAxML version 8: A tool for phylogenetic analysis and post-analysis of large phylogenies. Bioinformatics 30(9):1312–3.

81. Zankari E, Hasman H, Cosentino S, Vestergaard M, Rasmussen S, Lund O, Aarestrup FM, Larsen MV. 2012. Identification of acquired antimicrobial resistance genes. J Antimicrob Chemother 67(11):2640–4.

82. Davies TJ, Swann J, Sheppard AE, Pickford H, Lipworth S, Abuoun M, Ellington MJ, Fowler PW, Hopkins S, Hopkins KL, Crook DW, Peto TEA, Anjum MF, Walker AS, Stoesser N. 2023. Discordance between different bioinformatic methods for identifying resistance genes from short-read genomic data, with a focus on Escherichia coli. Microb Genomics 9(12):001151.

83. Edgar R, Bibi E. 1997. MdfA, an Escherichia coli multidrug resistance protein with an extraordinarily broad spectrum of drug recognition. J Bacteriol 179(7):2274–80.

84. Letunic I, Bork P. 2024. Interactive Tree of Life (iTOL) v6: Recent updates to the phylogenetic tree display and annotation tool. Nucleic Acids Res 52(W1):W78–82.

